# Entorhinal cortex signals dimensions of past experience that can be generalised in a novel environment

**DOI:** 10.1101/2025.08.01.668096

**Authors:** Sam Hall-McMaster, Lennart Wittkuhn, Luianta Verra, Noa L. Hedrich, Kazuki Irie, Peter Dayan, Samuel J. Gershman, Nicolas W. Schuck

**Affiliations:** Department of Psychology and Center for Brain Science, Harvard University; Max Planck Institute for Human Development; Institute of Psychology, University of Hamburg; Max Planck Institute for Biological Cybernetics; University of Tübingen; Max Planck UCL Center for Computational Psychiatry and Ageing Research

## Abstract

No two situations are identical. They can be similar in some aspects but different in others. This poses a key challenge when attempting to generalise our experience from one situation to another. How do we distinguish the aspects that transfer across situations from those that do not? One hypothesis is that the entorhinal cortex (EC) meets this challenge by forming factorised representations that allow for increased neural similarity between events that share generalisable features. We tested this hypothesis using functional magnetic resonance imaging. Female and male participants (n=40) were trained to report behavioural sequences based on an underlying graph structure. Participants then made decisions in a new environment where some, but not all graph transitions from the previous structure could be generalised. Behavioural results showed that participants distinguished the generalisable transition information. Accuracy was significantly higher in blocks where sequence transitions were shared across environments, than those in which transitions differed. This boost in accuracy was especially pronounced during early exposure to the novel environment. Throughout this early phase, neural patterns in EC showed a corresponding differentiation of the generalisable aspects. Neural patterns representing starting locations in familiar and novel environments were significantly more similar in EC on trials where sequences could be generalised from prior experience, compared to trials with new sequential transitions. This signalling was associated with improved performance when prior sequence knowledge could be reused. Our results suggest that during early exposure to novel environments, EC may signal dimensions of past experience that can be generalised.

**Significance Statement:** Generalisation is the process of using our experience to solve new challenges. A central problem we face when attempting to generalise is determining which aspects of a new situation are similar to what we have experienced before and which are different. This allows us to generalise our knowledge selectively, transferring insights to the aspects that are similar and relevant in novel scenarios, while avoiding over-generalisation to the aspects that differ. The results from this experiment suggest that the entorhinal cortex may distinguish these aspects during early exposure to new environments, supporting complex generalisation behaviour.

## Introduction

Generalisation is the process of using our experience to solve new challenges (Sharkey & Sharkey, 1993). One enduring view is that the way we generalise is guided by how similar two situations are in a “psychological space” (Shepard, 1987). What complicates this picture is that similarity can be multi-dimensional. Some information might be shared across situations but other information might not (Collins & Frank, 2013). Moreover, only some similarities may matter for our current goals. This raises a major question. How do our brains distinguish the dimensions relevant for generalisation in new situations or environments? As a simple example, imagine learning to drive on the right-hand side of the road and travelling to a country that drives on the left. While most knowledge about driving can be applied to the new context, the car position relative to oncoming traffic cannot. This unshared information is not just a neutral factor, but can actively interfere with the ability to perform well in new environments, a process known as negative transfer (Xia & Collins, 2021). To generalise effectively, therefore, one must be able to leverage shared aspects across situations whilst avoiding interference from other (non-shared) aspects.

Different areas within the medial temporal lobe (MTL) have been shown to distinguish shared and non-shared information across contexts, making them potential neural substrates for generalisation (Baram et al., 2024; Bhui et al., 2018; Shpektor et al., 2024; Whittington et al., 2020, 2022). Recent studies indicate that activity patterns in entorhinal cortex (EC) and hippocampus (HPC) are more similar across contexts with shared reward contingencies (Baram et al., 2021; Glitz et al., 2022) and are more distinct in HPC when stimuli are remapped to different positions in state space (Koolschijn et al. 2019).

The idea that EC and HPC distinguish shared information across contexts through changes in representational similarity is intriguing. However, it has yet to be tested in situations like the driving example, where environments are multi-dimensional and only some dimensions of earlier learning experiences can be generalised. To address the challenge of selecting dimensions for generalisation in new contexts or environments, one proposal is that the brain uses factorisation (Behrens et al. 2018; Whittington et al., 2020). Factorisation involves taking the different elements of an event or object and representing them separately, allowing particular dimensions to be selectively activated with minimal interference (Flesch et al., 2022; Lindsey & Issa, 2024) and for different dimensions to be combined in novel ways (Liu et al., 2019; Whittington et al., 2022). Factorised representations differ from conjunctive representations, which bind co-occurring dimensions into activity patterns that reflect the combination of features or elements (Komorowski et al., 2009; Rudy & O’Reilly, 2001). There is some evidence to suggest that spatial and object representations are factorised in EC (Behrens et al., 2018; Manns & Eichenbaum, 2006; Whittington et al., 2022) but conjunctive in HPC (Komorowski et al., 2009; Zheng et al., 2024). Based on these factors, one hypothesis is that EC contributes to generalisation by factorising events comprised of multiple dimensions, thereby allowing us to distinguish between generalisable and non-generalisable dimensions across environments (Behrens et al., 2018; Whittington et al., 2022).

Here we test the idea that factorised representations in EC play a role in rapidly determining which dimensions of past experience can be generalised in a novel environment. Participants (n=40) completed a planning task during functional magnetic resonance imaging (fMRI), in which transitions were based on a graph structure with multi-dimensional stimuli at each node. Participants completed the planning task in two distinct environments. Similar to the driving example, participants were extensively trained in one environment (the familiar environment). They were then placed in a new environment, where graph transitions were not known in advance (the novel environment). The task was designed so that the familiar and novel environments covertly shared graph transitions for one of the stimulus dimensions. If participants were able to factorise the stimuli and separate their dimensions, we expected they would then be able to reuse shared transitions from the familiar environment to enhance their performance in the novel environment.

Consistent with these ideas, we found that participants performed better on trials in the novel environment when transitions could be generalised from the familiar environment. In line with the hypothesis that EC would show a corresponding separation of the generalisable and non-generalisable dimensions, evident in levels of cross-environment representational similarity, we found that representational similarity in EC was significantly higher for stimulus features associated with generalisable transitions, during early exposure to the novel environment. These results point to a possible neural mechanism for distinguishing between aspects of past experience that can or cannot be reused, when we enter new environments and need to make decisions.

## Materials and Methods

### Participants

We set a target sample of 40 participants based on previous studies (Wittkuhn et al., 2021; Moneta et al., 2023). 52 participants were recruited to take part in the experiment, which involved three MRI scans lasting approximately one and half hours each. To progress with scans two and three, participants needed to complete a specific training protocol (described under pretraining for session two). Eight participants did not complete the training protocol in the time available and did not progress to scans two and three. One participant completed the training protocol but withdrew from the experiment at the start of scan two. Behavioural data from scan two was lost for one participant due to an error, in which the task was accidentally restarted and the original data file was overwritten. Two final participants were excluded due to strong head motion artefacts during the functional runs. Artefacts were evident during visual inspection of the images and converged with outliers identified in five MRIQC metrics (Esteban et al., 2017). The two participants had 12/32 and 15/32 functional runs with mean framewise displacements (Jenkinson et al., 2002) identified as outliers (more than three standard deviations above the sample mean), 6/32 and 8/32 runs with DVARS scores (Power et al., 2012) identified as outliers, 16/32 and 12/32 runs identified as outliers based on the AFNI outlier ratio, 8/32 and 12/32 runs identified as outliers based on the global signal correlation (Saad et al., 2013), as well as 11/32 and 9/32 runs identified as outliers based on low temporal SNR (Kruger et al., 2001). Exclusions due to data loss and motion did not influence the main generalisation effect seen in participant behaviour during scan three (final sample: n=40, M_diff_=7.68%, SD_diff_=17.56, permutation test: *p*=0.006, Fig. 5D; all available data: n=42, M_diff_=7.45%, SD_diff_=17.17, permutation test: *p*=0.005). The final sample included 40 participants, who were 18-34 (mean age=25, 27 female), had normal or corrected-to-normal vision (including normal colour vision) and reported no history of neurological or psychiatric illness. Participants who completed the full protocol received €110 and could earn up to €5 extra in each scan based on performance. The study was approved by the German Psychological Society and all participants signed informed consent before taking part.

### Materials

Stimulus presentation was controlled using Psychophysics Toolbox-3 (RRID: SCR_002881, version 3.0.17) in MATLAB (RRID: SCR_001622). Pretraining tasks were presented on a 15.6-inch screen with a spatial resolution of 1920 x 1080 and refresh rate of 60Hz outside the scanner (MATLAB version 2014b). Responses were recorded with D, F, J, K and L keys on a standard QWERTY keyboard. Scanning tasks were run on a stimulus computer with a spatial resolution of 1024 x 768 and a refresh rate of 16.7Hz, which projected to a magnetically safe screen for participants inside the scanner (MATLAB version 2017b). Responses were recorded using two fiber optic button boxes (Current Designs, Philadelphia, PA, USA). Participants used two buttons on the left box (middle finger and index finger) and three buttons on the right box (index finger, middle finger and ring finger). MRI data was preprocessed using ReproIn (RRID: SCR_017184, HeuDiConv version 0.9.0), fMRIprep (Esteban et al., 2019, RRID: SCR_016216, version 20.2.4), with FSL (Jenkinson et al., 2012, RRID: SCR_002823, version 1.3.3) and MRIQC (Esteban et al., 2017, version 0.16.1) being used to assess image quality. Behavioural and MRI analyses were performed in Python (RRID:SCR_008394, version 3.8.19).

### Data and Code Availability

Data and code materials will be made available on publication.

### Experimental Tasks

Participants performed a series of zookeeper-themed learning and decision-making tasks. The tasks used ten animal stimuli, divided into two zoo sections (north and south). Participants first learned that each animal was associated with a specific food pellet. Food pellets were unique combinations of shapes and textures. The same five shapes and the same five textures were used in the two zoo sections (north and south), but were combined in different ways for each animal. After learning the associations, participants learned that animal enclosures in the north section of the zoo were structured in a particular way. The structure meant that certain animal enclosures would always be encountered after other enclosures when travelling in the zoo. Based on this structure, participants were taught to plan sequences of four food pellets that would be needed for a trip around the north section of the zoo, from each starting enclosure. To assess generalisation, participants were tested on their ability to plan trips in the south section of the zoo, despite not being taught about the arrangement of these enclosures. Unbeknownst to participants, there was a hidden structure that could be reused across zoo sections. Sequences for one food feature (e.g. shapes) were actually the same in both zoo sections. Sequences for the other feature did not share any of the same pairwise transitions (see Fig. 1). Analyses focused on whether participants showed higher accuracy on blocks that used generalisable information than non-generalisable information.

**Figure 1.**
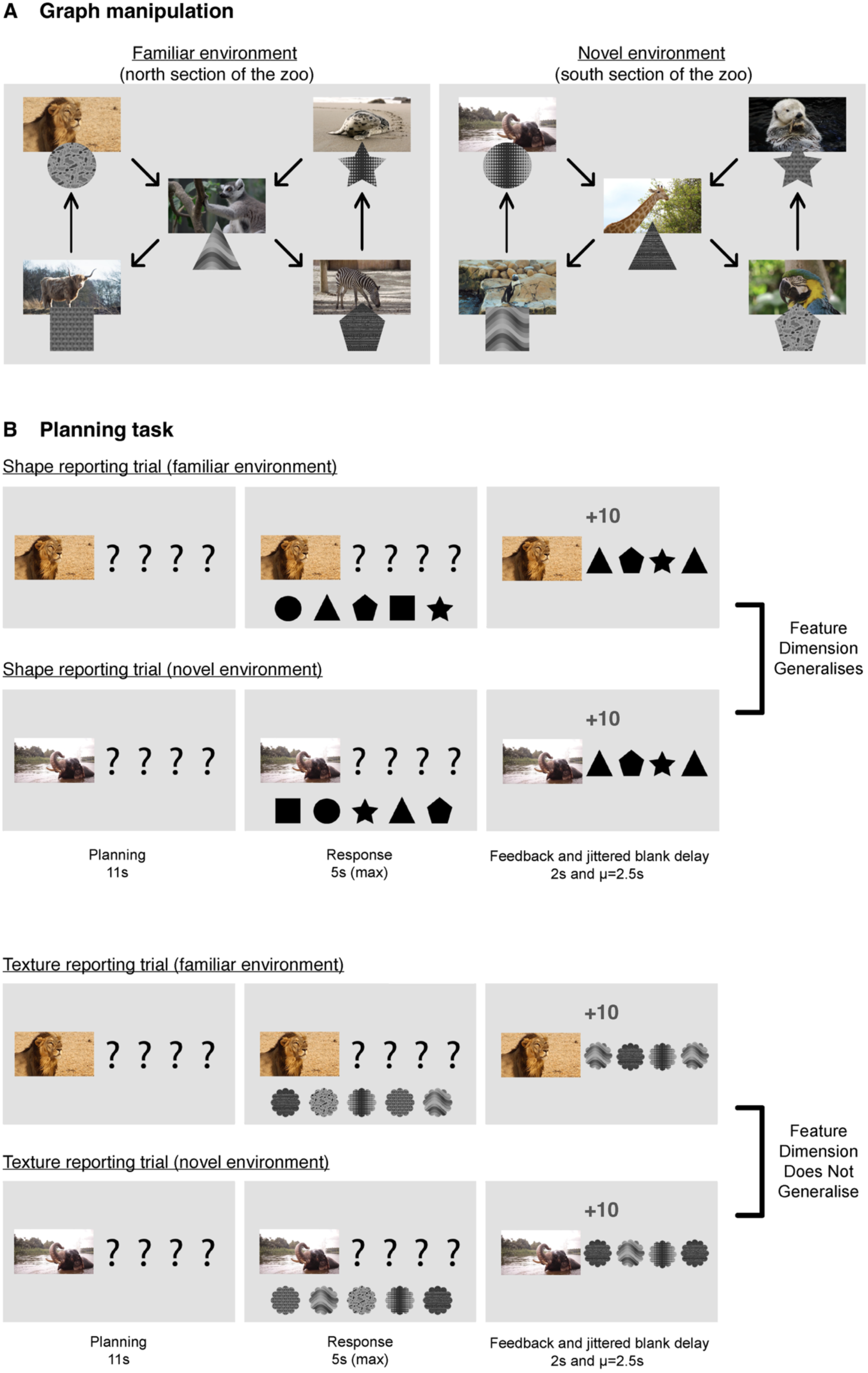
Experimental design. **A:** Participants completed a planning task while being scanned with fMRI. The task involved two different environments defined as graphs. Participants received extensive pre-training on the graph transition structure for one environment before scanning (the familiar environment) but were not pre-trained on transitions for the other environment before scanning (the novel environment). The experiment was designed so that some transition information was secretly shared across environments. For some participants, shape transitions were preserved across graphs and the texture transitions differed (as in the example above). For the other participants, the opposite was true. This created an experimental setting in which participants needed to distinguish the shared and unshared feature dimensions across contexts for successful performance in the novel environment. **B:** The cover story was that each participant was a zookeeper responsible for feeding the animals in a virtual zoo. On each trial, participants were shown an animal cue that indicated their current location within the zoo. Participants then needed to plan an ordered sequence of four food items (textured shapes) necessary for feeding the animals on a trip around the zoo. Participants were given 11s to plan and then 5s to enter their intended sequence during a response phase. Stimuli available for selection during the response phase were presented in random screen positions on every trial and only after the planning phase, to ensure sequences of food items had no systematic relationship with the motor sequences used to report them. A subsequent feedback screen showed the participants entered sequence (if correct) or an example of a correct sequence (if incorrect). Some blocks of the planning task asked participants to report shape sequences and in others asked them to reported texture sequences. For each participant, one feature dimension could be generalised across environments (e.g. shapes). This meant that from a given starting location (e.g. the top left graph node), the same stimulus sequence would be correct in both environments. The other feature dimension could not be generalised (e.g. textures). This meant that from a given starting location (e.g. the top left graph node), different sequences were required in each environment for correct performance.

#### Session one (localiser)

##### Pretraining one

In the session one pretraining, participants learned that each animal was associated with a specific food pellet. Food pellets were bi-dimensional stimuli composed of a shape and a texture. On each trial, participants were shown a short animal video (0.75s), followed by a 0.5s blank delay. The five food pellets from the current zoo section were then shown on screen in a random order. Participants needed to press the key (D, F, J, K, or L) that corresponded to the screen position of the correct food pellet within 5s. Following a response, feedback was shown for 2s (‘Correct!’ or ‘Incorrect!’). If the response was incorrect, participants were shown the correct animal-food pellet pairing. The trial concluded with a blank inter-trial interval (ITI) lasting 1s. Participants began the pretraining task in a randomly determined section of the zoo, continuing with blocks of 21 trials in that section until they had scored at least 19/21 trials correct in a block (90%). Participants then progressed to learning about the other zoo section. After passing the 90% criterion for both zoo sections, participants had to do so one more time but with a shorter response deadline of 1.75s. Pairwise stimulus transitions across trials were balanced in each block, ensuring that each animal was followed by all other animals in its zoo section the same number of times.

##### Scan one

The scan started with a resting state functional measurement, in which they were asked to fixate on a centrally presented cross for three minutes. Participants then performed the task from pretraining task 1. Participants were shown a brief animal video and asked to imagine its corresponding food pellet before selecting it from a response array within 1.75s. The setup was similar to the pretraining, except for the timing and numbers of trials/blocks. The blank delay following the animal video was drawn from a truncated exponential distribution, with a mean of 3s (lower bound=1s, upper bound=12s). Feedback (+1 or +0) was 0.5s. The ITI was drawn from a truncated exponential distribution with a mean of 1.5s (lower bound=0.5s, upper bound=12s). Participants completed 61 trials per block, with fully balanced pairwise transitions between animals across trial pairs. Participants began in a random section of the zoo and alternated zoo section from block-to-block. After four task blocks, fieldmaps and a five-minute T1w anatomical measurement were made. Participants then completed another four blocks. This resulted in 488 trials, 48 trials for each of the ten animal stimuli (excluding the first trial from each block which was selected to balance the pairwise transitions). Scan one concluded with a three-minute resting state measurement. To motivate good performance, participants were shown their average accuracy and median RT at the end of each block. Participants could also unlock bonus items (e.g. a cartoon guitar) for every 12.5% of the maximum points earned on the task.

#### Session two (familiar environment)

##### Pretraining two

In the pretraining for session two, participants were taught about the sequential structure of the animals in the north section of the zoo. Sections were structured so that certain animal enclosures would only be encountered after visiting other enclosures. The graph structure of the zoo sections is shown in Figure 1. Pretraining two consisted of two main parts. In the first part, participants learned pairwise transitions between the animal enclosures. On each trial, participants were shown an animal video for 0.75s, with the question “Which animal comes next?” printed above it. Following this, the animal video stayed on the final frame as a still image and two alternative animal images were shown in the bottom left and bottom right corners of the screen. One of the animals shown followed the correct sequence and the other was a distractor. Whether each of the target animals appeared on the left or right side of the screen was selected at random. Participants had up to 2 seconds to choose an animal. If the choice was correct, participants saw the cue animal and the selected animal with “Correct!” printed above. If the choice was incorrect, participants saw the cue animal and the target animal with “Incorrect! The correct order was:” printed above. Feedback was displayed for 2s. The trial concluded with a 1s ITI. At the beginning of the task, participants did not know the pairwise transitions and had to learn through trial and error. Participants completed six trials per block, one for each starting animal, plus an additional trial for the animal in the middle of the graph which could transition to two possible subsequent enclosures. Participants needed to score 100% on a block to continue to the next phase. Note that participants only completed training 2 for one section of the zoo (north).

Following pairwise learning, participants transitioned to learning about the main planning task in preparation for scan two. This task showed participants at a starting point in the zoo (i.e. a starting animal) and asked them to imagine the next food pellet they would need on a trip around the zoo. The intention was that participants imagined the subsequent animal in the sequence and retrieved its associated food pellet. On each trial, participants were shown a starting animal (0.75) and had 4s to imagine the upcoming pellet that was needed. Participants then saw the possible food pellets on screen, presented in a random order. Participants needed to select the position of the correct pellet. This response format meant it was only possible to imagine the identity of the upcoming pellet in the 4s interval before the response phase. The motor response required to select that item could not be determined in advance, before the possible response options were shown. If the response was correct, “Correct!” was printed to the screen. If the response was incorrect or too slow, “One possible sequence was:” was printed to the screen with an example of a correct sequence. Feedback lasted 2s, followed by a 1s ITI. Each block contained 12 trials, two for each starting position, plus two extra trials for the middle node in the graph that had two possible subsequent enclosures. On trials involving the middle node, the next animal in the sequence was presented on screen during the planning phase as a still image, so that participants had practice planning along both branches of the graph. Participants needed to get one correct response for each trial type (minimum 6/12) to advance to the next stage.

Upon advancing, participants needed to plan the next two food pellets. The planning phase and response limit both increased to 6s and participants needed to reach the same criteria to advance. The training continued to build up the planning length to 4 items. This was met with corresponding increases in trial duration (8s for planning and 8s to respond for 3 items; 10s for planning and 8s to respond for 4 items). Once participants could successfully plan four food pellets, different blocks were introduced based on sensory feature. In some blocks, participants needed to plan four food shapes from the starting location. In other blocks, they needed to plan four food textures. The number of trials per block was reduced to 10 because participants were able take a branch of their choosing from the middle node. This branching structure at the middle node meant that, from this point on, there was more than one valid path from each starting location. Participants needed to get one correct response from each starting animal to pass the block and had to do for one shape and one texture block to advance. The final stage was identical except that the response phase was reduced to 5s in preparation for the scanning task. The short response interval was designed so participants needed to prepare as much as possible during the planning phase.

##### Scan two

In scan two, participants completed the final task from pretraining two. The scan began with a three-minute resting state measurement. Whether participants started with a shape or texture planning block was randomly determined. The feature used for planning then alternated from block-to-block. On each trial, participants were shown an animal video (0.75s) and had 11s to plan the subsequent four food features needed on a trip around the zoo. The response items were then presented in a random order on screen and participants needed to select the positions of the four items in the correct sequence. This response format meant that sequence of food features had no systematic relationship with the motor sequences used to report them. This was designed to isolate the planning of stimulus sequences during planning phase, by making participants unable to determine the button presses sequence needed to implement their plan until later in the trial. Participants completed 20 trials per block, with four trials for each starting position. Trial order was randomised. From the fifth participant onwards, the randomisation was constrained so that the same animal cue could not appear two trials in a row. After four blocks, a three-minute resting state measurement was taken, followed by fieldmaps and a five-minute T1w measurement. Participants completed an additional four blocks and the scan concluded with a three-minute resting state measurement. As in scan one, participants could unlock bonus items for every 12.5% of the maximum points earned on the task.

#### Session three (novel environment)

##### Pretraining three

To prepare for scan three, participants completed the same task as in pretraining one for the south section of the zoo. This zoo section did not appear in pretraining two or scan two and pretraining three was used as a reminder of the animal-food pellet pairs in this section. Participants needed to score 19/21 (90%) on one block with a response deadline of 1.75s to complete pretraining three.

##### Scan three

In scan three, participants completed the same scanning protocol as scan two, except with the animals for the south section of the zoo. Participants had not been trained on the graph structure for these animals and needed to learn their arrangement during the scan itself. Unbeknownst to participants, we created a shared structure across scan two and scan three. Specifically, the sequential order of one feature (shape or texture) was actually the same in both scans. The sequential order of the remaining feature was different and did not share any of the same pairwise transitions as scan two. This meant that sequential knowledge about one feature was generalisable across scans, while the other was non-generalisable. Whether shape or texture could be generalised was approximately balanced across participants (shape = 19 participants, texture = 21 participants). Whether the first block in scan three required the generalisable or non-generalisable feature for planning was also approximately balanced (participants for whom shape was generalisable: began with shape = 9, began with texture = 10; participants for whom texture was generalisable: began with texture = 10, began with shape = 11). As in scans 1 and 2, participants could unlock bonus items for every 12.5% of the maximum points earned on the task.

##### Debrief

To assess participants’ knowledge about the task structure, the session concluded with a debrief task outside the scanner. Participants were asked whether they had noticed that the sequences of shapes or the sequences of textures had repeated across scans 2 and 3 and gave a binary response (yes or no). This was followed by a question asking, if they had to guess, which feature was repeated across the scans (shape or texture).

### Behavioural Analyses

Task performance was assessed on the basis of transition accuracy. Each trial required a valid 4-item sequence from the cued starting to location to receive reward. Transition accuracy was calculated per trial as the proportion of consecutive sequence items in a valid order. Transitions accuracies were averaged across trials and compared for blocks in the novel environment where prior sequences could or could not be reused. Statistical comparisons were made using two-sided permutation tests with 10,000 permutations to generate the null distributions. We predicted that participants would show significantly higher accuracy in the novel environment on blocks where sequences could be reused from the familiar environment.

#### Baseline performance

Knowledge of the animal-object associations during the localiser scan, and planning accuracy in the familiar environment were assessed with one-sample permutation tests. The level of performance expected due to chance was subtracted from each participant score and the sample mean was calculated. On each permutation, the sign was randomly flipped for approximately half of the participant scores and the sample mean was recomputed and stored in the null distribution. This repeated for 10,000 permutations. The true sample mean was then compared against the 95^th^ percentile of the null distribution to determine significance. A paired permutation test was used to compare planning accuracy in the familiar environment for the feature that would and would not become generalisable later in the experiment. Each permutation involved randomly flipping the condition label for approximately half the participants. As a sanity check, we further evaluated whether participants exhibited a significant bias for using the left or right branch at the hub node in the familiar environment. To do so, we calculated the difference in the number left and right transitions entered from the hub during the response phase for each participant. The resulting values were converted to proportions (-1=complete rightward bias, +1=complete leftward bias). The participant distribution was then tested against 0 using a one-sample permutation test. Resulting *p*-values for the tests in this section were corrected using the Benjamini-Hochberg correction (Benjamini & Hochberg, 1995).

#### Generalisation performance

Behavioural analyses focused on average generalisation performance are described at the beginning of Behavioural Analyses section. To build on these and account for trial-level accuracy, we used a linear mixed effects model. Trial accuracy was the dependent variable. Predictors included feature generalisability (whether transitions be generalised on the current trial), trial index (log-transformed to account for non-linear changes in accuracy over time), starting position within the graph, block type (whether participants needed to report shape or texture sequences) and cumulative exposure to each of the six possible node transitions based on presented feedback up to that point. The model included random intercepts for each participant and was implemented using statsmodels (www.statsmodels.org).

#### Impact of awareness

To assess whether explicit awareness of the generalisable feature dimension influenced accuracy in the novel environment, we used a linear mixed effect model. The model included awareness as a between-subjects factor (aware vs. unaware), as well as block type (generalisable vs. non-generalisable) and phase (early vs. late) as within-subject factors. Random intercepts and slopes were used for each within-subject factor for each participant. Likelihood ratio tests were then used to compare the full model to reduced models omitting each fixed effect. Resulting *p*-values were corrected for seven tests (three main effects and four interaction tests) using the Benjamini-Hochberg correction (Benjamini & Hochberg, 1995).

#### Learning analyses

To better understand how participants discovered and applied the generalisable feature dimension, we quantified reuse of familiar transitions in the novel environment, and examined how this changed over time. For each trial in the novel environment, we summed the number of valid pairwise transitions entered during the response phase. The sum was converted to a proportion by dividing it by four, the maximum number of transitions per trial. We then tested whether the average reuse proportion on the first trial in the novel environment was significantly different from chance. To do so, we separated participants into two groups: those who started with a generalisable block in the final scanning session and those who did not. The two distributions were then separately tested against chance. This procedure ensured we were examining reuse on the very first trial in the generalisable environment for each block type. To examine broad directional trends in reuse for each condition, we fit separate linear regression models predicting the proportion of reuse based on trial number. To examine the contributions of seeing particular transitions during feedback, we conducted regression analyses that tested whether the transitions entered on each trial could be predicted by (a) transitions shown during feedback on the previous trial and (b) the cumulative number times each transition had been shown as feedback up that point. For the non-generalisable condition, we additionally made use of the familiar environment stimulus space during scoring. This allowed us to ask whether seeing a novel transition as feedback increased or decreased the likelihood of using a corresponding transition from the familiar environment on the current trial. For example, if one saw a novel transition (e.g. square followed by circle) as feedback, we could ask whether a corresponding familiar transition (e.g. square followed by triangle) was more or less likely to be entered on the current trial. Distributions of beta coefficients were compared against 0 using one-sample permutation tests. Resulting *p*-values were corrected using the Benjamini-Hochberg correction (Benjamini & Hochberg, 1995).

### MRI parameters

Data were acquired using a 32-channel head coil on a 3-Tesla Siemens Magnetom TrioTim MRI scanner (Siemens, Erlangen, Germany). Functional measurements were whole brain T2* weighted echo-planar images (EPI) with multi-band (MB) acceleration (voxel size=2×2×2×mm; TR=1250ms; echo time (TE)=26ms; flip angle (FA)=71 degrees; 64 slices; matrix = 96 x 96; FOV = 192 x 192mm; anterior-posterior phase encoding direction; distance factor=0%; MB acceleration=4). Slices were tilted +20° (counter-clockwise) relative to the corpus callosum, with the aim of improving signal quality from hippocampus (Weiskopf et al., 2006). Functional measurements began after five TRs (6.25s) to allow the scanner to reach equilibrium and help avoid partial saturation effects. 145 volumes were acquired during each resting state run. Up to 377 volumes (scan one) or 341 volumes (scans two-three) were acquired during each task run; acquisition was ended earlier if participants had completed all trials. Fieldmaps were measured using the same scan parameters, except that two short runs of 20 volumes were collected with opposite phase encoding directions.

Fieldmaps were later used for distortion correction in fMRIPrep (Esteban et al., 2019, RRID: SCR_016216, version 20.2.4). Anatomical measurements were acquired using T1 weighted Magnetization Prepared Rapid Gradient Echo (MPRAGE) sequences (voxel size=1×1×1×mm; TR=1900ms; TE=2.52ms; flip angle FA=9 degrees; inversion time (TI)=900ms; 256 slices; matrix = 192 x 256; FOV = 192 x 256mm).

### MRI Preprocessing

Preprocessing was performed using fMRIPrep 20.2.4 (Esteban, Markiewicz, et al. (2018); Esteban, Blair, et al. (2018); RRID:SCR_016216), which is based on Nipype 1.6.1 (Gorgolewski et al. (2011); Gorgolewski et al. (2018); RRID:SCR_002502).

#### Anatomical Data Preprocessing

A total of 3 T1-weighted (T1w) images were found within the input BIDS dataset. All of them were corrected for intensity non-uniformity (INU) with N4BiasFieldCorrection (Tustison et al. 2010), distributed with ANTs 2.3.3 (Avants et al. 2008, RRID:SCR_004757). The T1w-reference was then skull-stripped with a Nipype implementation of the antsBrainExtraction.sh workflow (from ANTs), using OASIS30ANTs as target template. Brain tissue segmentation of cerebrospinal fluid (CSF), white-matter (WM) and gray-matter (GM) was performed on the brain-extracted T1w using fast (FSL 5.0.9, RRID:SCR_002823, Zhang, Brady, and Smith 2001). A T1w-reference map was computed after registration of 3 T1w images (after INU-correction) using mri_robust_template (FreeSurfer 6.0.1, Reuter, Rosas, and Fischl 2010). Brain surfaces were reconstructed using recon-all (FreeSurfer 6.0.1, RRID:SCR_001847, Dale, Fischl, and Sereno 1999), and the brain mask estimated previously was refined with a custom variation of the method to reconcile ANTs-derived and FreeSurfer-derived segmentations of the cortical gray-matter of Mindboggle (RRID:SCR_002438, Klein et al. 2017). Volume-based spatial normalization to two standard spaces (MNI152NLin6Asym, MNI152NLin2009cAsym) was performed through nonlinear registration with antsRegistration (ANTs 2.3.3), using brain-extracted versions of both T1w reference and the T1w template. The following templates were selected for spatial normalization: FSL’s MNI ICBM 152 non-linear 6th Generation Asymmetric Average Brain Stereotaxic Registration Model [Evans et al. (2012), RRID:SCR_002823; TemplateFlow ID: MNI152NLin6Asym], ICBM 152 Nonlinear Asymmetrical template version 2009c [Fonov et al. (2009), RRID:SCR_008796; TemplateFlow ID: MNI152NLin2009cAsym].

#### Functional Data Preprocessing

For each of the 32 BOLD runs found per subject (across all tasks and sessions), the following preprocessing was performed. First, a reference volume and its skull-stripped version were generated using a custom methodology of fMRIPrep. A B0-nonuniformity map (or fieldmap) was estimated based on two (or more) echo-planar imaging (EPI) references with opposing phase-encoding directions, with 3dQwarp Cox and Hyde (1997) (AFNI 20160207). Based on the estimated susceptibility distortion, a corrected EPI (echo-planar imaging) reference was calculated for a more accurate co-registration with the anatomical reference. The BOLD reference was then co-registered to the T1w reference using bbregister (FreeSurfer) which implements boundary-based registration (Greve and Fischl 2009). Co-registration was configured with six degrees of freedom. Head-motion parameters with respect to the BOLD reference (transformation matrices, and six corresponding rotation and translation parameters) are estimated before any spatiotemporal filtering using mcflirt (FSL 5.0.9, Jenkinson et al. 2002). BOLD runs were slice-time corrected to 0.588s (0.5 of slice acquisition range 0s-1.18s) using 3dTshift from AFNI 20160207 (Cox and Hyde 1997, RRID:SCR_005927). The BOLD time-series were resampled onto the following surfaces (FreeSurfer reconstruction nomenclature): fsnative. The BOLD time-series (including slice-timing correction when applied) were resampled onto their original, native space by applying a single, composite transform to correct for head-motion and susceptibility distortions. These resampled BOLD time-series will be referred to as preprocessed BOLD in original space, or just preprocessed BOLD. The BOLD time-series were resampled into standard space, generating a preprocessed BOLD run in MNI152NLin6Asym space. First, a reference volume and its skull-stripped version were generated using a custom methodology of fMRIPrep. Several confounding time-series were calculated based on the preprocessed BOLD: framewise displacement (FD), DVARS and three region-wise global signals. FD was computed using two formulations following Power (absolute sum of relative motions, Power et al. (2014)) and Jenkinson (relative root mean square displacement between affines, Jenkinson et al. (2002)). FD and DVARS are calculated for each functional run, both using their implementations in Nipype (following the definitions by Power et al. 2014). The three global signals are extracted within the CSF, the WM, and the whole-brain masks. Additionally, a set of physiological regressors were extracted to allow for component-based noise correction (CompCor, Behzadi et al. 2007). Principal components are estimated after high-pass filtering the preprocessed BOLD time-series (using a discrete cosine filter with 128s cut-off) for the two CompCor variants: temporal (tCompCor) and anatomical (aCompCor). tCompCor components are then calculated from the top 2% variable voxels within the brain mask. For aCompCor, three probabilistic masks (CSF, WM and combined CSF+WM) are generated in anatomical space. The implementation differs from that of Behzadi et al. in that instead of eroding the masks by 2 pixels on BOLD space, the aCompCor masks are subtracted a mask of pixels that likely contain a volume fraction of GM. This mask is obtained by dilating a GM mask extracted from the FreeSurfer’s aseg segmentation, and it ensures components are not extracted from voxels containing a minimal fraction of GM. Finally, these masks are resampled into BOLD space and binarized by thresholding at 0.99 (as in the original implementation). Components are also calculated separately within the WM and CSF masks. For each CompCor decomposition, the k components with the largest singular values are retained, such that the retained components’ time series are sufficient to explain 50 percent of variance across the nuisance mask (CSF, WM, combined, or temporal). The remaining components are dropped from consideration. The head-motion estimates calculated in the correction step were also placed within the corresponding confounds file. The confound time series derived from head motion estimates and global signals were expanded with the inclusion of temporal derivatives and quadratic terms for each (Satterthwaite et al. 2013). Frames that exceeded a threshold of 0.5 mm FD or 1.5 standardised DVARS were annotated as motion outliers. All resamplings can be performed with a single interpolation step by composing all the pertinent transformations (i.e. head-motion transform matrices, susceptibility distortion correction when available, and co-registrations to anatomical and output spaces). Gridded (volumetric) resamplings were performed using antsApplyTransforms (ANTs), configured with Lanczos interpolation to minimize the smoothing effects of other kernels (Lanczos 1964). Non-gridded (surface) resamplings were performed using mri_vol2surf (FreeSurfer). Many internal operations of fMRIPrep use Nilearn 0.6.2 (Abraham et al. 2014, RRID:SCR_001362), mostly within the functional processing workflow. For more details of the pipeline, see the section corresponding to workflows in fMRIPrep’s documentation.

### Neural Analyses

Bilateral anatomical masks for EC, HPC were generated for each participant using FreeSurfer cortical parcellations. For pipeline validation, bilateral masks were generated for occipital cortex. For exploratory analyses, bilateral masks were generated for medial orbitofrontal cortex, rostral anterior cingulate cortex, precuneus, interior parietal cortex, and the rostral middle frontal gyrus as a proxy for dorsolateral prefrontal cortex. For more information on this aspect, please see Desikan et al. (2006) and the FreeSurfer documentation: https://surfer.nmr.mgh.harvard.edu/fswiki/FsTutorial/AnatomicalROI/FreeSurferColorLUT

To empirically test whether EC distinguished information that could be generalised from the familiar to the novel environment, we used representational similarity analysis (RSA; Kriegeskorte et al., 2008). RSA is useful because – unlike standard decoding approaches that focus on one variable at a time - it allows one to assess the contribution of multiple variables to the neural signal simultaneously. This in turn allows one to: a) compare the contribution between variables based on a single analysis (e.g. assessing how much the neural signal changes as a function of shared vs. unshared information across environments) and b) control for variables that are expected to contribute to the neural signal but are not of immediate interest (e.g. differences in performance between familiar and novel environments). RSA has been successfully used in range of previous studies on sequence learning and decision-making (e.g. Bellmund et al., 2022; Brunec & Momennejad, 2022; Hall-McMaster et al., 2022; Myers et al., 2015; Luyckx et al., 2019). To apply RSA to our dataset, we undertook the following steps. For each ROI, we extracted the voxel activity from the onset of the animal cue, on each trial in scans 2 and 3. Data were spatially smoothed by 5mm (Baram et al., 2024; Shpektor et al., 2024). The resulting data were detrended, z-scored and motion confounds (identified by fMRIPrep) were removed. Data cleaning was performed separately for each functional run using Nilearn signal clean. The data were then temporally smoothed to stabilise the neural patterns by averaging the neural signal at each TR with the TR before and after it. Trial data was subdivided into conditions based on the environment (familiar/scan 2 vs. novel/scan3), block type (shape or texture reporting block) and the cued starting position (graph node 1-5). Neural similarity between each condition pair was estimated with cross-validated Mahalanobis distances (crossnobis distances), which give unbiased similarity estimates in the presence of noise (Schütt et al., 2023; Walther et al., 2016). The crossnobis distance between two conditions (A, B) is defined as:

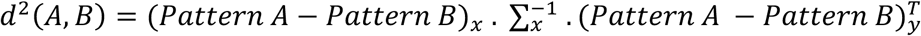

The first term is the difference in mean voxel patterns between the two conditions from one half of the data (partition x). The second term is the inverse of an error covariance matrix, estimated from one half of the data (partition x). Calculation of the error covariance matrix used a shrinkage estimator to down-weight noisy covariance estimates (Ledoit & Wolf, 2004). The final term is the transpose of the difference in the mean voxel patterns between the two conditions from the remaining half of the data (partition y). Unless otherwise stated, all trials for each condition were used in the crossnobis distance calculation. The calculation was performed in a time-resolved manner in that we extracted activation patterns across trials at a common TR from the starting point of the planning phase (e.g. the third TR from cue onset). Data centered on that TR was used as input to the crossnobis distance calculation. The same procedure was then applied to data centered on other TRs (the fourth TR from cue onset, the fifth TR and so on). For each participant-ROI combination, this resulted in a 20×20 representational dissimilarity matrix (RDM) for each TR, a total of 16 TRs from the start of the planning phase (i.e. 0-20s from cue onset). We next constructed model RDMs to reflect expected neural structure based on different task variables of interest. The logic of all models was to place zeroes in cells of a 20 × 20 matrix where conditions matched on the variable of interest and ones in all remaining cells. For example, the ‘animal model’ had zeroes in each cell of a matrix where two conditions involved the same animal cue (e.g. lion). The upper triangular portion of each matrix was then transformed into a vector and z-scored. The resulting data and model distance vectors were then entered into a multiple regression analysis testing that tested whether model vectors were significant predictors of the neural vector at each TR. This resulted in one *β* coefficient per model RDM per measurement volume from cue onset. We were primarily interested in understanding representational similarity during the planning phase. We therefore averaged each *β* coefficient of interest from +3.75 to +15s following cue onset, assuming an approximate hemodynamic delay of 4s. The resulting participant distribution was tested against 0 using two-sided permutation tests with 10000 permutations to generate the null distribution. The null distribution was generated by randomly flipping the sign of each participant coefficient in 10000 permutations and computing the mean coefficient for the sample. If the true sample mean fell within the 95^th^ percentile of the null distribution, the neural similarity was deemed significant.

#### Validation

To validate our RSA pipeline, we examined whether information about the animal cues could be recovered from occipital cortex (OC), as we expected this region to show strong, differentiated responses to different visual stimuli. The anatomical mask used for this ROI combined FreeSurfer parcellations for the cuneus, lateral OC and pericalcarine regions. To ensure stable neural RDMs, crossnobis distances were computed 25 times based on random trial splits for each condition. The average across splits was used for the final RDM. Our validation analysis used the following general linear model and was run separately at each TR. Note that neural RDMs were computed using all trials (unless otherwise stated) and each TR here refers to a common timepoint across trials from planning phase onset. DV stands for distance vector.

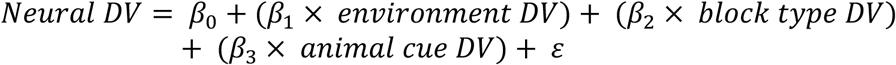

The resulting *β*_3_coefficients for the animal cue were averaged from +3.75 to +15s following cue onset for each participant. The participant distribution was tested against 0 using a two-sided permutation test. A significant above chance result was taken as evidence for significant neural coding of the animal cues and validation of the RSA pipeline.

#### Neural signalling of generalisable feature information

To empirically test whether EC distinguished information that could be generalised between the familiar and novel environments, we constructed two critical model RDMs (see Fig. 2). These had 0s in the cells comparing conditions across the familiar and novel environment where the same stimulus feature was cued by the presented animal (e.g. a circle). The “gen. model” RDM had this structure for condition comparisons where sequences could be generalised across environments. The “non-gen. model” had this structure for condition comparisons where sequences could not be generalised across environments. We tested whether these model RDMs were significant predictors of the neural data from EC and HPC. The crossnobis distance calculation used to generate neural RDMs for EC and HPC involves randomly splitting the trials from each condition into two partitions. To ensure stability, we ran this process 250 times and took the average distance estimate for each cell in the neural RDMs.

**Figure 2.**
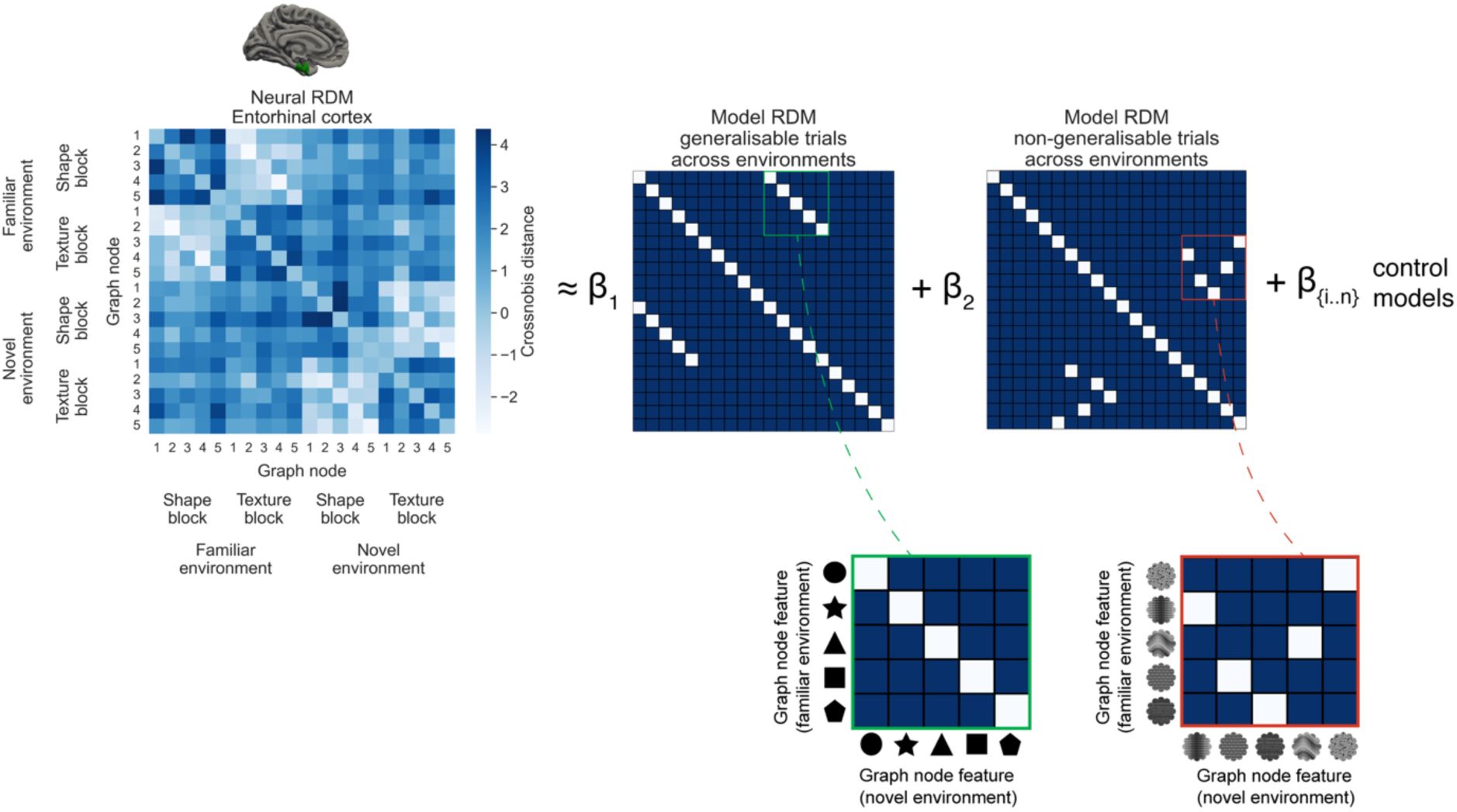
Neural analysis framework. To test whether MTL subregions distinguish generalisable and non-generalisable information across environments, we performed a representational similarity analysis (RSA). We first extracted neural data from the entorhinal cortex (EC) and computed cross-validated Mahalanobis distances (crossnobis distances) between all condition pairs. Unlike metrics that are not cross-validated, crossnobis distances can take on negative values when patterns contain noise. If the true distance is 0, this will result in distance estimates symmetrically distributed around 0 (Schütt et al., 2023). We then asked whether the resulting representational dissimilarity matrix (RDM) exhibited a relative increase in neural similarity for stimuli at the starting location on each trial when transition information could be generalised across environments. The “gen. model” RDM (shown beside *β*_1_) was formulated to capture cross-environment stimulus similarity for the generalisable feature dimension. In this case, each stimulus occupied the same graph node in both environments and therefore sequential transitions for this dimension could be generalised (shown in the green inset linked to the *β*_1_ RDM). The “non-gen. model” RDM (shown beside *β*_2_) was formulated to captured cross-environment stimulus similarity for the feature dimension that did not generalise. In this case, each stimulus was mapped to different node positions across graphs and therefore transitions for this feature dimension differed in the two environments (shown in the red inset linked to the *β*_2_ RDM). These model RDMs were used as predictors in a multiple regression to explain the neural RDM. The regression procedure simultaneously controlled for a variety of other task factors that could plausibly influence the neural signal, including differences in accuracy between conditions and differences in visual information shown on screen. As a final step, we tested whether *β*_1_ was significantly higher than *β*_2_ as a representational marker distinguishing generalisable from non-generalisable trials across environments. The same process was also applied to neural data extracted from the hippocampus (HPC).

In addition to the gen. and non-gen. model RDMs, we constructed control RDMs to account for other aspects of the neural signal (Fig. 3). These included models for condition differences in the environment, block type, animal cue, as well as individualised control RDMs for the number of on-screen stimulus matches during the response and feedback phases, and condition differences in accuracy on the current and previous trial. Binary accuracy (the proportion of fully correct sequences reported) was used for the accuracy model RDMs, as opposed to transition accuracy (the proportion of correct transitions reported). This was because we were interested in testing whether neural signals distinguishing generalisable information had a functional connection to performance (transition accuracy) in the novel environment. Removing covariance from the neural signal associated with transition accuracy as a control RDM would be expected to eliminate a potential association between these variables. Using binary accuracy allowed us to control for overarching condition differences in accuracy, while still retaining relevant covariation for neuro-behavioural tests. In addition to the control model described so far, we formulated control RDMs to account for various graph properties. These included model RDMs for the shortest bidirectional distance between nodes (link distance), the average bidirectional distance between nodes, the one-step bidirectional transitions within each environment-block context, and the node type (i.e. whether a node was a hub in the graph, an input to the hub or an output from the hub). The upper triangular portion of the neural and model RDMs were transformed into vectors and z-scored. The subsequent distance vectors were then entered into the following general linear model at each TR. Neural RDMs were computed using all trials (unless otherwise stated) and each TR here refers to a common timepoint across trials from the start of the planning phase.

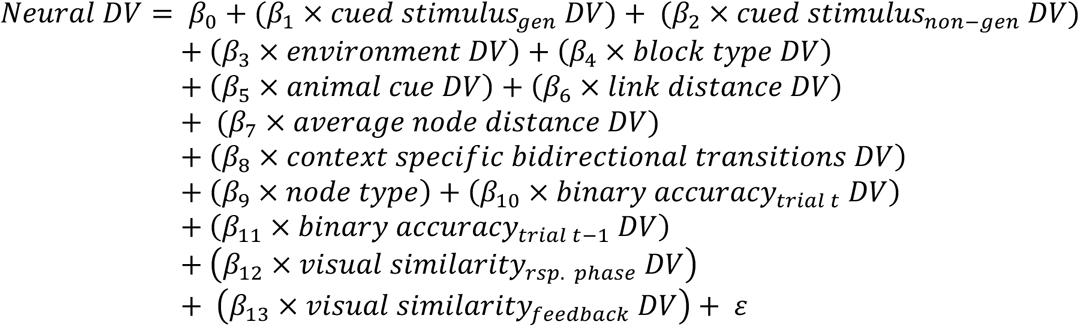

**Figure 3.**
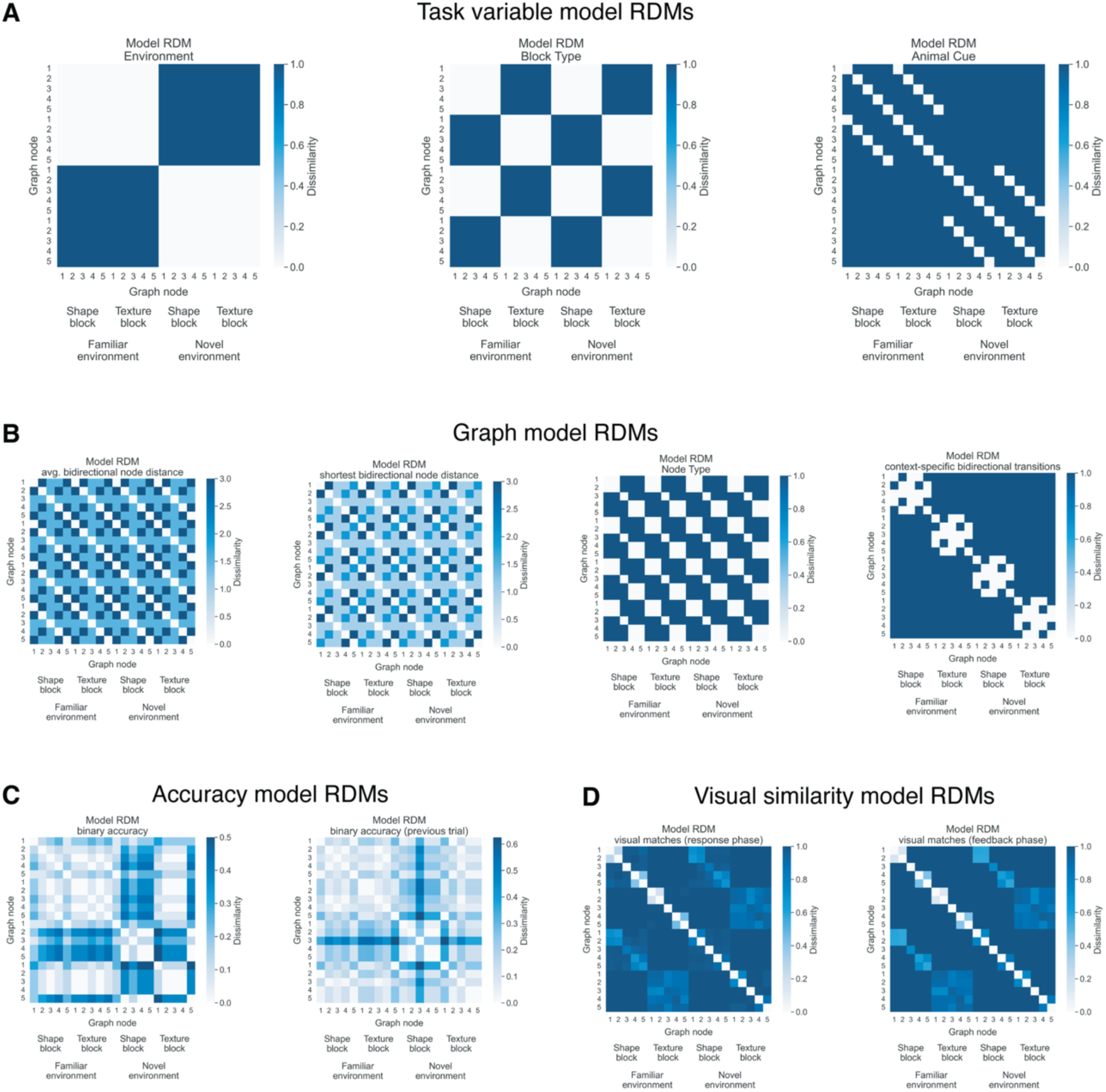
Control representational dissimilarity matrices (RDMs). **A:** RDMs capturing condition differences for the main task variables. These include environment (novel vs. familiar), block type (report shape sequences vs. report texture sequences) and animal cue. **B:** RDMs capturing condition differences based on the graph structure. These include model RDMs for the average bidirectional distance between nodes, the shortest bidirectional distance between nodes (i.e. the link distance), the node type (i.e. whether a node was a hub in the graph, an input to the hub or an output from the hub) and one-step bidirectional transitions within each environment-block context. **C:** Example model RDMs capturing differences in binary accuracy between conditions for an individual participant. This measure is the difference in proportion of fully correct sequence entries between two conditions. Model RDMs capture differences in accuracy based on the current and previous trial, respectively. **D:** Example model RDMs capturing how often stimuli presented on-screen match between conditions for an individual participant. Model RDMs capture such visual matches during the response and feedback phases, respectively. **A-D:** Lighter colours indicate a relative increase in similarity for all RDMs.

The resulting *β*_1_and *β*_2_ coefficients were averaged from +3.75 to 15s following cue onset. To test whether representational similarity across environments was significantly above zero in each ROI, we compared the resulting *β*_1_and *β*_2_coefficients against zero using two-sided permutation tests. To assess whether MTL subregions were distinguishing shared and unshared elements across environments, we tested whether the *β*_1_ and *β*_2_ coefficients were significantly different from each other using two-sided paired permutation tests. Condition labels for the coefficients were randomly swapped over 10000 iterations and the mean coefficient difference between conditions was computed to generate the null distribution. If the true sample difference fell within the 95^th^ percentile of the null distribution, neural similarity was deemed significantly different in the generalisable and non-generalisable blocks of the novel environment. To test for a neural distinction during early exposure to the novel environment, we repeated the analysis pipeline but restricted the analysis to only use trials from the first half of scanning in the novel environment (trials 1-80 of a possible 160) to compute crossnobis distance estimates. Coefficient tests against 0 were corrected for multiple comparisons using the Benjamini-Hochberg correction (Benjamini & Hochberg, 1995). Tests of the difference between generalisation coefficients (*β*_1_ vs. *β*_2_) were uncorrected because they were derived from prior theoretical and empirical work (Baram et al., 2021; Behrens et al. 2018; Glitz et al., 2022; Koolschijn et al. 2019; Whittington et al., 2022).

#### Neural signalling of generalisable transition information

To test whether the neural distinction between generalisable and non-generalisable feature dimensions extended past the cued staring location in each graph, we repeated the previous analysis but included three additional model RDMs for one-step forward transitions (Fig. 4). The models assumed that when an animal cue was shown in the novel environment, the next feature would be reactivated. Like the previous section, we used cross-environment similarity to isolate specific feature transitions (e.g. a specific shape independent from the texture and animal cue). Within-environment comparisons can be used to assess transition information in the neural signal but only for combinations of animal cues/features. Two model RDMs were used to isolate one-step feature transitions in the generalisable blocks and non-generalisable blocks in response to each cue in the novel environment. A final model RDM was used to isolate feature transitions in the non-generalisable blocks based on the old transition structure from the familiar environment, which was no longer relevant. The resulting coefficients were compared in a 3×2 repeated measures ANOVA, with factors of transition model (gen. forward vs. non-gen. forward vs. non-gen. irrel. forward) and ROI (EC. vs. HPC).

**Figure 4.**
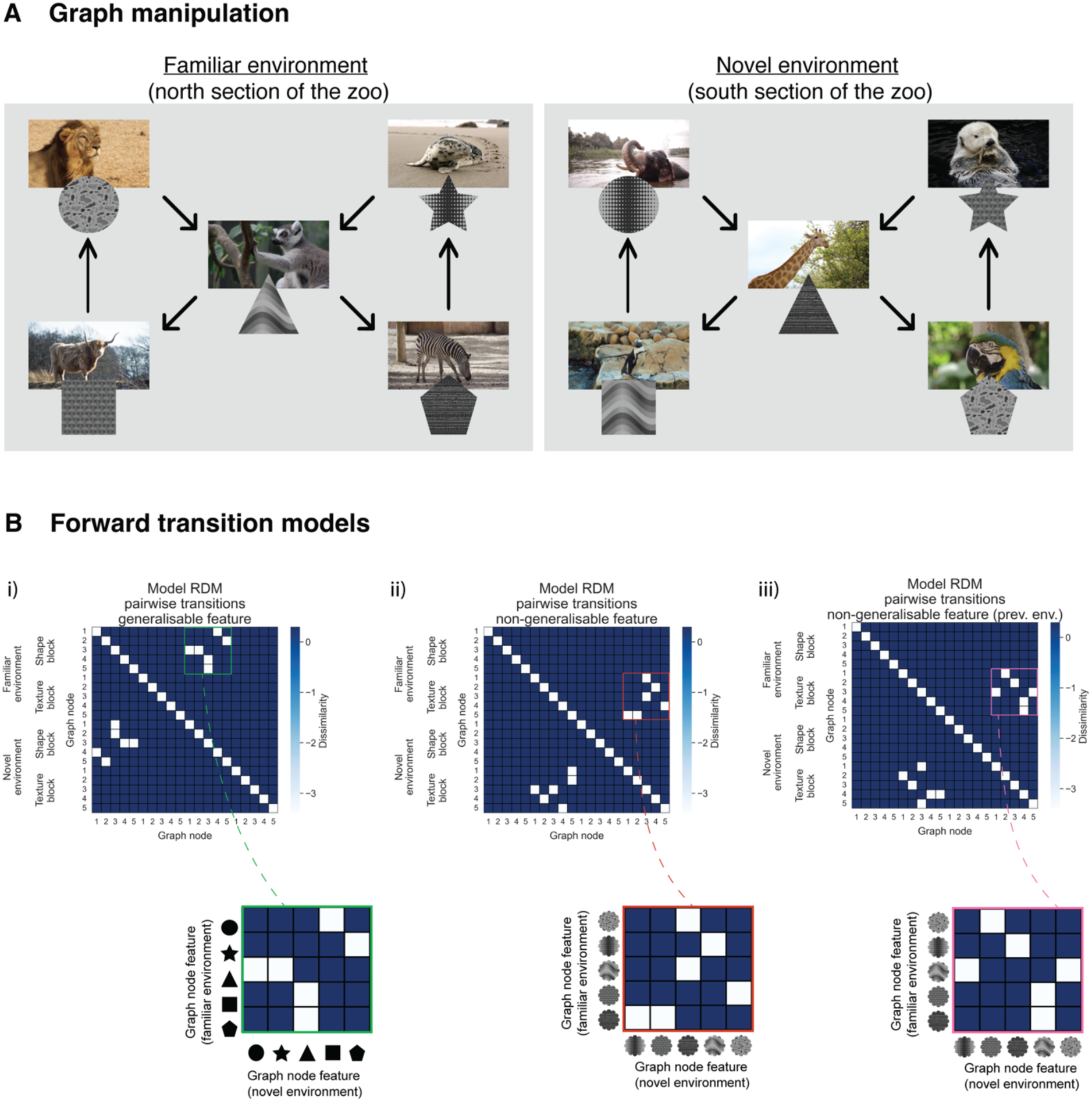
Model representational dissimilarity matrices (RDMs) for one-step forward transitions. **A:** Example graphs for an individual participant, as shown in the main text. **B:** Example forward transition models. The main aim of the models is to test the idea that when an animal cue is shown in the novel environment, the stimulus at the next node in the graph gets activated. This activation is then expected to increase the neural similarity between the current condition in the novel environment and the condition in the familiar environment that began with that activated feature. **i):** The model RDM on the left captures one-step forward transitions for the generalisable blocks of the novel environment. For example, seeing an elephant cue in the novel environment during a shape block (the circle node), would lead to the activation of the triangle as the next node, increasing the similarity with lemur trials in the familiar environment where the triangle was cued. **ii):** The model RDM in the middle captures one-step forward transitions for the non-generalisable blocks. For example, seeing an elephant cue in the novel environment during a texture block (the node with the dotted-column texture), would lead to the activation of the horizontal line texture as the next node, increasing the similarity with zebra trials in the familiar environment where that texture was cued. **iii):** The model RDM on the right captures one-step forward transitions in the non-generalisable blocks, based on the irrelevant graph structure from the familiar environment. For example, seeing an elephant cue in the novel environment during a texture block (the node with the dotted-column texture), it would be as if the participant was in the top right node of the familiar environment (a node which also has the dotted-column texture). Planning one step from this point would lead to activation of the wave texture, increasing the similarity with lemur trials from the familiar environment.

#### Neural signalling and performance

Spearman correlations were used to test whether neural signals identified in the EC were correlated with performance in the novel environment. The variables used in the correlations were the difference between the gen. and non-gen. model coefficients (*β*_1_and *β*_2_) in EC based on trials 1-80 in the novel environment (1-80) and the difference in performance on those same trials between generalisable or non-generalisable blocks. Follow-up tests examined whether the neural difference correlated with performance in the generalisable or non-generalisable blocks individually. A Williams test was used to examine whether the correlation strength was significantly higher for the neuro-behavioural correlation involving generalisable blocks compared to the correlation involving non-generalisable blocks. The Benjamini-Hochberg procedure was used to correct for equivalent correlation tests across ROIs (Benjamini & Hochberg, 1995).

## Results

To test the hypothesis that EC distinguishes dimensions of past experience that can be generalised in novel environments, 40 participants completed an fMRI experiment in which they navigated non-spatial “zoo” environments (Fig. 1). Each environment was defined as a graph. Each node in the graph corresponded to an animal enclosure and an associated “food” stimulus, where food stimuli were textured shapes (e.g. a triangle filled with dots, a striped circle etc). In a baseline session, participants learned the animal-food pairings for all graph nodes (e.g. that lions were fed circular speckled food etc). Participants were then pre-trained on graph transitions from the “north” section of the zoo, learning deterministic paths among five animal enclosures and their associated food stimuli. During scanning, participants completed a planning task that required them to traverse the north section and feed the animals from cued starting points, reporting the sequence of food shapes or food textures needed along the route. On each trial, participants saw an animal cue that indicated their starting location and had a 11s to plan an ordered sequence of four food stimuli from the starting point. The graph structure permitted more than one valid path from each starting location and each animal could be visited up to two times within a path. After the planning phase, a set of response options appeared on screen in a random order and participants had up to 5s to enter their plan, by selecting the screen position of each sequence item in turn. The random ordering of response options meant that planned sequences of food stimuli had no systematic relationship with the motor sequences needed to enter them across trials. Because the trial specific order was only presented after the planning phase, the decoupling also prevented participants from preparing a sequence of motor responses or button presses during the planning phase. Following the response phase, participants saw a feedback screen showing their planned sequence (if correct) or an example of a correct sequence (if incorrect). In a subsequent session, participants were scanned again while learning to navigate the “south” section of the zoo. This environment was novel to participants because it contained different zoo animals from the previous environment and participants were not taught transitions between graph nodes in advance. Unbeknownst to them, the novel environment secretly contained a mixture of shared and unshared food sequence transitions with the familiar environment. Specifically, the novel south section contained the same food shapes and textures as before but in new combinations (e.g. a striped triangle). This allowed us to transfer sequential transitions for one feature dimension (e.g. shapes) across the familiar and novel zoo environments, while introducing new sequential transitions for the other feature (e.g. textures). As a consequence, reusing multi-dimensional transitions from the familiar environment in the form of conjunctive representations would hinder performance in the novel environment, because half of the transitions no longer applied. In contrast, factorising representations and only considering transitions along the shared feature dimension would help performance.

### Baseline performance

Participants learned the animal-object associations for both zoo sections to high degree (M_acc_=96.18%, SD_acc_=3.56, chance=20%, permutation test: *p*<0.001, Fig. 5A), as assessed during a localiser scan. Planning accuracy was calculated per trial as the proportion of consecutive sequence items in a valid order. Participants showed high planning accuracy in the familiar environment during scanning (M_acc_=93.77%, SD_acc_=4.73, chance=24%, permutation test: *p*<0.001, Fig. 5B), for which they received extensive pre-training. Within this baseline environment, we did not detect significant differences in accuracy between the feature sequences that would ultimately become generalisable or non-generalisable later in the experiment (M_gen_=93.44% vs. M_non-gen_=94.09%, M_diff_=-0.66%, SD_diff_=3.85, permutation test: *p*=0.419, Bayes Factor favouring the null hypothesis=5.05, Fig. 5C). The graph structure included a central hub node from which participants could freely decide between two valid transitions (e.g. the lemur in Fig 1A). Exploratory analyses of the baseline environment did not detect a significant group level bias in the percentage of choices using the left or right branch at the hub node (M=-0.82%, SD=67.83, *p=*0.940). While left/right choices were unbiased across participants, they tended to be consistent within participants: examining branch choices at the individual participant level showed that 31 of the 40 participants had 95% confidence intervals that did not overlap with 50%, suggesting a reliable preference for using one branch over another in those individuals. Most crucially, the analyses in this section suggest that there was no pre-existing difference in performance between generalisation conditions, before participants were exposed to the novel environment.

**Figure 5.**
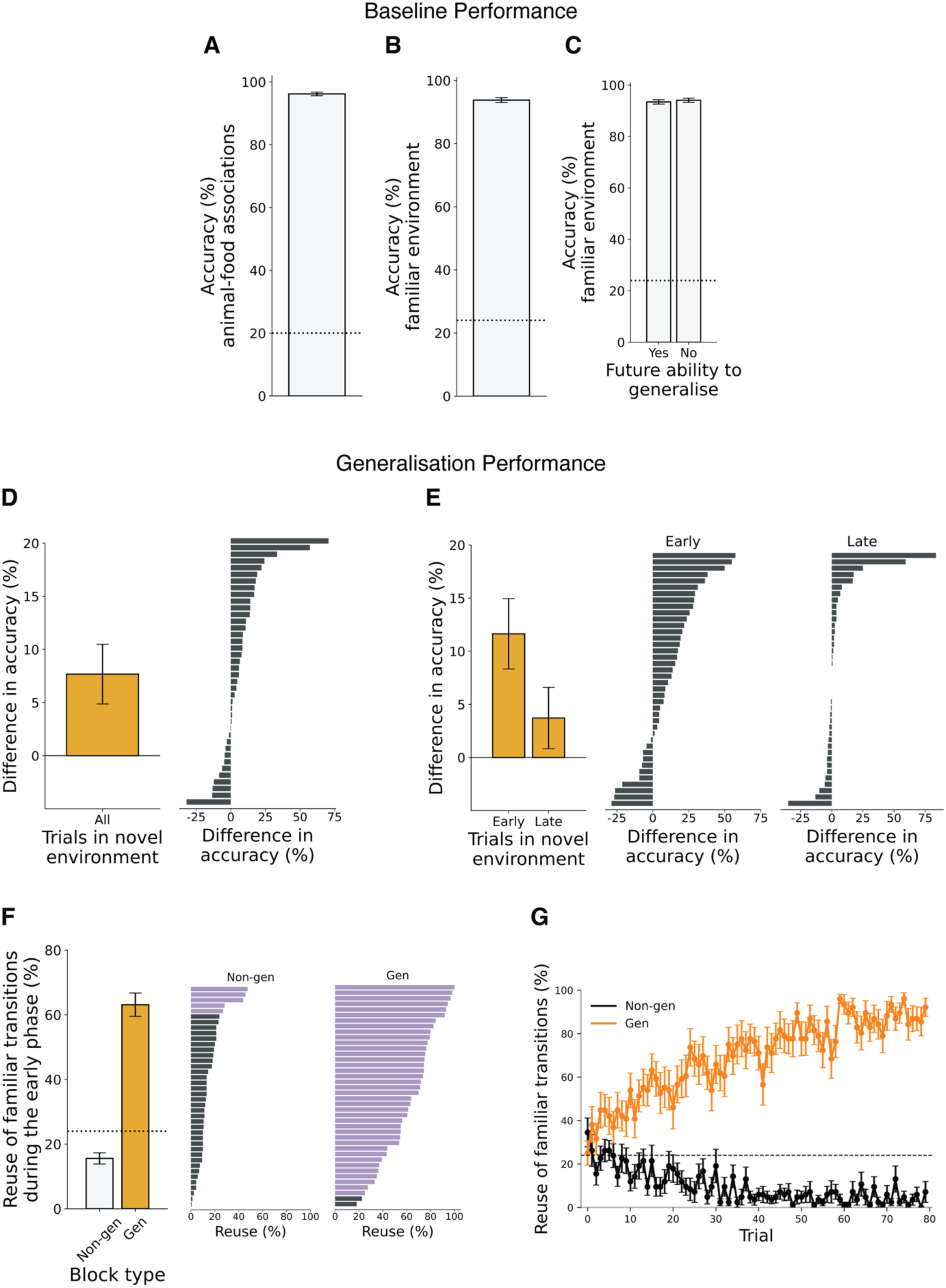
Behavioural performance. **A:** Accuracy in retrieving the relevant food target (textured shape) when shown each animal cue during a localiser MRI scan. **B:** Accuracy during planning trials in the familiar environment during a second MRI scan. **C**: Accuracy in B separated based on whether sequences could be generalised to the novel environment later in the experiment. A-C: The dotted line indicates chance. **D:** The difference in accuracy between blocks in the novel environment where previously learned sequences could be generalised and blocks where previous sequences did not generalise during a third MRI scan. Higher values indicate greater relative accuracy on blocks with generalisable information. The corresponding tornado plot shows difference scores for individual participants. **E:** The accuracy difference in D separated into early trials (1-80) and late trials (81-160) in the novel environment. Tornado plots show difference scores for individual participants. **F:** How often transitions from the familiar environment were reused during early trials (1-80) in the novel environment. The measure is expressed as a percentage of response opportunities. The plot is separated into blocks where participants needed to plan based on the generalisable or the non-generalisable feature dimension. The dotted line indicates chance. The corresponding tornado plots show scores for individual participants. Scores in purple are numerically above chance and scores in black are numerically below chance. **G:** Reuse of familiar transitions in the novel environment across trials. Different lines are used to show reuse for trials where transitions generalise or do not generalise across environments. To ensure the reuse curves reflect performance from the very first trial in the novel environment, participants were separated into two groups: those who started with a block where the active feature dimension generalised and those who started with a block where the active feature dimension did not generalise. Each line therefore reflects data from half of the sample. The dotted line indicates chance. A-G: In all plots, errors bars show the standard error of mean.

### Participants generalised prior learning

To test for generalisation in the novel zoo environment, we examined whether accuracy was higher in blocks where sequences from the familiar environment could be reused. We expected that reusable information would confer a performance benefit in the novel environment and hence that accuracy would be higher for blocks where transitions could be reused. Consistent with this idea, accuracy was significantly higher in task blocks where transitions generalised across environments, compared to blocks where transitions differed (M_gen_=68.91% vs. M_non-gen_=61.23%, M_diff_=7.68%, SD_diff_=17.56, permutation test: *p*=0.006, Fig. 5D). This performance benefit for shared feature information was especially pronounced during the first half of scanning in the novel environment (M_gen_=55.72% vs. M_non-gen_=44.08%, M_diff_=11.64%, SD_diff_=20.68, permutation test: *p*<0.001, Fig. 5E). In the second half of scanning in the novel environment, performance was not significantly different across feature conditions (M_gen_=82.11% vs. M_non-gen_=78.39%, M_diff_=3.72%, SD_diff_=18.05, permutation test: *p*=0.248), representing a significant attenuation in the performance difference compared to the early phase (M_early vs. late_=7.92%, SD_early vs. late_=16.56, permutation test: *p*=0.004, Fig. 5E). Linear mixed effects (LME) modelling of trial-level accuracy showed consistent results. The model included predictors of feature generalisability, trial index (log-transformed to account for non-linear learning dynamics), starting node, block type (shape vs. texture), cumulative exposure to each transition based on presented feedback, and random intercepts for each participant. The model showed a significant main effect of generalisability on trial accuracy (*β*=0.24, SE=0.04, 95% CI [0.17 0.32], *z*=6.19, *p*<0.001), indicating higher performance on trials where transitions from the familiar environment could be generalised. The model also showed a significant negative interaction between feature generalisability and the log-transformed trial index (*β*=−0.04, SE=0.01, 95% CI [−0.06, −0.02], *z*=−4.40, *p*<0.001), indicating the relative performance boost on generalisable trials attenuated with time, as participants gained more experience in the novel environment.

One way the performance difference between conditions could emerge during the early phase would be if participants attempted to unselectively generalise all transitions from familiar environment to the novel environment. This would boost performance in the generalisable blocks but impair performance in the non-generalisable blocks. We found that the performance difference in this early stage was not due to over-generalisation of transitions from the familiar environment. To the contrary, participants showed a below chance tendency to enter familiar transitions in the novel environment during blocks that contained new sequential transitions (M=15.58%, SD=10.84, chance=24%, permutation test: *p*=0.002, Fig. 5F). For example, if shape transitions could be generalised but texture transitions could not, participants were less likely than chance to repeat a texture transition from the familiar environment during the texture reporting blocks in the novel environment. Using the graphs from Fig. 1 to illustrate, a participant starting at the elephant/dotted texture in a texture reporting block of the novel environment would be less likely than chance to select the wave texture, even though the dotted texture is followed by the wave texture in the familiar environment. Hence, the performance difference above reflected adaptive use of generalisable transitions in one stimulus dimension (e.g. shape), but not the other (e.g. texture). Further supporting the idea that reuse was adaptively calibrated, we found that lower reuse on blocks with new sequential transitions during the early phase was associated with significantly higher accuracy on blocks with generalisable transitions (Spearman’s *ρ*(38)=0.56, *p*<0.001). To understand whether individual differences in path preference in the familiar environment were related to broad path preferences in the novel environment, we examined whether the tendency to repeatedly use one branch at the hub node in the familiar environment was correlated with an equivalent tendency to repeatedly use one branch in the novel environment. Exploratory Spearman correlations revealed that the degree of branch preference in the familiar environment was significantly related to the degree of branch preference in non-generalisable trials (*ρ*(38)=0.38, *p*=0.021), but not generalisable trials (*ρ*(38)=0.22, *p*=0.182), during the early phase in the novel environment.

We next examined whether explicit awareness of the shared structure influenced performance benefits. A post-experimental questionnaire indicated that 27.5% of participants noticed the repetition and correctly identified the feature with preserved sequential structure. To test whether this awareness increased the difference in performance between generalisable and non-generalisable blocks in the novel environment, we used a LME model. The model included awareness as a between-subjects factor (aware vs. unaware), as well as block type (generalisable vs. non-generalisable) and phase (early vs. late) as within-subject factors. Random intercepts and slopes were used for each within-subject factor for each participant. Likelihood ratio tests were then used to compare the full model to reduced models omitting each fixed effect. As expected from our earlier results testing the prediction of improved performance for generalisable information, the model indicated significant effects of block type (*β*=0.13, SE=0.04, 95% CIs=[0.20 0.05], χ²(1)=13.24, *p*=0.036) and phase (*β*=0.25, SE=0.04, 95% CIs=[0.17 0.32], χ²(1)=106.68, *p*<0.001) on accuracy in the novel environment. However, awareness (*β*=0.05, SE=0.09, 95% CIs=[0.23 - 0.13], χ²(1)=1.83, *p*=0.938) and interactions that included awareness were not significant (*β*s<0.07, χ²(1) values<1.43, *p*-values>0.568). This suggests that having explicit awareness of which feature transitions generalised did not have a significant influence on performance in the novel environment.

In a final set of behavioural analyses, we sought to understand more about how the generalisable feature was discovered and applied. We considered two high-level strategies that one could employ in the novel environment. First, one could generalise both features from the outset. If this were true, reuse of transitions from the familiar environment should be significantly above chance for both features on the first trial in the novel environment. Second, one could start out by making random choices in the novel environment. If this were true, reuse of transitions from the familiar environment should not significantly differ from chance for either feature on the first trial in the novel environment. From either baseline (above chance or at chance), reuse should then further increase or decrease as a function of feature generalisability. Recall that participants were shown an example of a correct sequence during the feedback phase if their response was incorrect on a trial. On average, responses on trial 1 in the novel environment were not significantly different from the chance rate expected for random choice, both for participants who could generalise a previous transition on trial 1 (due to starting with a gen. block) and for participants who could not generalise on trial 1 (due to starting with an non-gen. block; M_gen_=25%, SD_gen_=24.33, chance=0.24, permutation test: *p*=0.819; M_non-gen_=34.52%, SD_non-gen_=30.35, chance=0.24 permutation test: *p*=0.118, Fig. 5G). From these starting points, trial number was a significant positive predictor of reuse on trials with generalisable transitions (*β*=0.007, SD=0.004, permutation test: *p*<0.001) and a significant negative predictor of reuse in early blocks with novel transitions (*β*=-0.002, SD=0.002, permutation test: *p*<0.001). This implies that participants started out in the novel environment with behaviour that was not significantly different from chance. However, adaptive reuse was subsequently calibrated based on increasing experience and feedback. To understand how feedback was related to increased or decreased reuse, we conducted regression analyses that examined potential predictors of using specific transitions in the novel environment. Starting with the generalisable blocks, we asked whether the specific transitions entered on the current trial could be predicted by (a) the transitions presented during feedback on the previous trial and (b) how many times each transition had been presented as feedback in total up to that point. Both variables were significant predictors of the transitions entered during trials with shared transitions across environments (*β*_prev. fbk_=0.04, SD=0.08, permutation test: p=0.006; *β*_total. fbk_=0.18, SD=0.10, permutation test: p<0.001). For non-generalisable blocks in the novel environment, we asked whether entering transitions from the familiar environment on the current trial was negatively predicted by the feedback variables (a) and (b), since the feedback in these blocks involved new sequential transitions. This revealed that cumulative presentation of each novel transition during feedback predicted decreased reuse of corresponding familiar transitions (*β*_total. fbk_=-0.20, SD=0.10, permutation test: *p*<0.001). To take an example from Fig. 1, the more one saw the transition from the dotted texture (elephant node) to the horizontal line texture (giraffe node) during feedback, the less likely one would be to use the dotted texture/wave texture transition from the familiar environment on the current trial. For non-generalisable blocks, feedback on the previous trial was not a significant predictor of transitions entered on the current trial (*β*_prev. fbk_=-0.01, SD=0.05, permutation test: *p*=0.137). These findings suggest that specific transitions shown during trial feedback contributed to increasing reuse of familiar transitions in the generalisable blocks and decreasing reuse in the non-generalisable blocks.

Overall, our behavioural results indicate that performance in the novel environment was better when information from the familiar environment could be reused, compared to cases where information about sequential transitions needed to be learned anew. This effect was most prominent during early exposure to the new environment and was not explained by a pre-existing difference in performance.

### Neural analysis framework

Having shown that decision performance in a new environment differed for sequences that could or could not be generalised from a previous environment, we sought to test whether the MTL tracked this distinction. To do so, we performed a representational similarity analysis (RSA; Kriegeskorte et al., 2008). The main components of this framework are shown in Figure 2. fMRI measurements from each participant were separated into conditions based on the environment (familiar vs. novel), block type (report shapes vs. report textures) and starting position in the graph (node 1-5). For each condition, the corresponding neural pattern was defined as the temporally smoothed voxel-wise activation pattern from three regions of interest (ROIs): primary occipital cortex (used for validation), EC and HPC. Activation patterns were extracted separately at each measurement timepoint starting from planning phase onset (time shifted by 3 TRs/+3.75s to account for hemodynamic lag). Each pattern was then temporally averaged with a sliding window (previous, current and next TR) to increase stability. Neural similarity between each condition pair was computed using cross-validated Mahalanobis distances (crossnobis distances; Schütt et al., 2023; Walther et al., 2016). When comparing each condition pair (e.g. lion and elephant trials across environments), we used all trials associated with the two conditions. Exceptions to this are stated in specific result sections. From these trials, we extracted temporally averaged data from a common timepoint, locked to the starting point of the planning phase (e.g. the third TR after cue onset). The resulting data was used to estimate the crossnobis distance for that condition pair at that TR during the planning phase. We then repeated this process for each common measurement volume from planning phase onset (the fourth TR, the fifth TR etc.), each pair of conditions and ROI. For each participant-ROI combination, this resulted in one representational dissimilarity matrix (RDM) per timepoint from planning phase onset. Since neural RDMs were computed across multiple trials, each timepoint here refers to the TR common across trials (e.g. TR_+3_ from the start point of the planning phase). We then constructed three model RDMs to capture possible task factors driving neural similarity. The model RDMs were matrices with binary entries that encoded whether a given variable matched for each pair of conditions. Matches were encoded with 0s to indicate that neural patterns associated with the two conditions were expected to have lower dissimilarity scores. Non-matches were encoded with 1s to indicate that neural patterns were expected to have higher dissimilarity scores. One model captured whether a pair of conditions was from the same environment (familiar vs. novel), another model captured whether a pair of conditions was from the same block type (report shapes vs. report textures) and another captured whether the animal cue presented on screen was the identical or different (Fig. 3). To relate the model RDMs to the neural RDM, we performed multiple regression at each timepoint, using the model RDMs to predict neural dissimilarity between conditions. The resulting beta coefficients were averaged over the 11s planning phase. Within this framework, higher coefficients suggest stronger neural coding of a given task variable.

To test whether this approach was robust, we performed a validation analysis using data from primary occipital cortex (OC). Specifically, we asked whether trials which involved the same animal would show higher similarity/lower dissimilarity, as we would expect this region to show distinct neural responses for different visual cues. The animal cue RDM was a significant predictor of the neural RDM from OC (mean *β*_animal_=0.43, SD=0.08, permutation test: *p*<0.001) indicating that our analysis pipeline could successfully recover expected information from the task.

### Entorhinal cortex signalled generalisable information in a novel environment

Having validated the analysis pipeline, we turned to our attention to EC and HPC. Past research has implicated both the EC and HPC in distinguishing shared and unshared information across well-learned or explicitly instructed contexts (Baram et al., 2021; Glitz et al., 2022; Mark et al., 2024; Koolschijn et al. 2019). However, the EC in particular has been proposed to represent information with factorised neural codes that allow different dimensions to be flexibility used for generalisation (Behrens et al., 2018; Whittington et al., 2020). Based on these ideas, we tested whether the EC exhibited a relative increase in neural similarity across environments when feature sequences could be generalised, consistent with a factorised coding scheme that allows feature dimensions to be separated and selectively activated (Fig. 2).

Testing this required a wider suite of model RDMs. We therefore constructed two additional model RDMs to measure the cross-environment similarity of neural representations (Fig. 2). The “gen. model” focused on comparing conditions with shared sequences across the familiar and novel environments. An example (used for participants who could generalise shape transitions) is shown in Fig. 2 next to *β*_1_ and includes a green inset that highlights the important cross-environment comparisons. The gen. model RDM was ultimately a matrix with 0s in the cells comparing conditions that started with the same relevant feature in both environments and allowed for the same sequences to be used from that starting point. For example, the lion and elephant in Fig. 1 are both associated with a circle and shape stimuli are positioned in the same graph nodes across environments. The gen. model RDM would therefore encode the comparison between lion and elephant trials during shape reporting blocks as a 0. Other cells in the matrix that did not meet these criteria had 1s to indicate higher expected neural dissimilarity scores. The “non-gen. model” followed similar logic except that it focused on cross-environment comparisons where sequences did not generalise. These were cases where the same relevant feature was cued at the starting location but different sequences were then required. For example, the lemur and the penguin in Fig. 1 are both associated with the wave texture but different sequences are needed each animal because texture stimuli are remapped across environments. The non-gen. model would encode the comparison between lemur and penguin trials during texture reporting blocks as a 0, and other comparisons that did not meet its criteria as 1s. An example non-gen. model (used for a specific participant as non-gen. features were remapped to across environments independently for each person) is shown in Fig. 2 next to *β*_2_, coefficients were averaged from coefficients were averaged from coefficients were averaged from inset that highlights the important cross-environment comparisons. Using these models as predictors for the neural RDMs in EC and HPC allowed us to test whether a) similar neural codes were used to represent the cued stimulus features across familiar and novel environments and b) whether the degree of neural similarity distinguished trials where sequences could be generalised or not. The stimuli displayed on screen in comparisons across environments were non-identical for both the gen. and non-gen. models. This means that differences in similarity based on these models should reflect differences in internal representations for the generalisable and non-generalisable dimensions.

In addition to the gen. and non-gen. model RDMs, we included a comprehensive set of control RDMs in our analysis procedure to account for other variables that could influence neural activity in EC and HPC (Fig. 3). This set included model RDMs for the environment, block type and animal cue used in the validation analysis. It also included model RDMs to control for various graph properties, including the link distance and average bidirectional distance between abstract graph nodes, bidirectional one-step transitions specific to each environment-block type combination, and whether the starting node was a hub in the graph, an input to the hub or an output from the hub. In comparison to the gen. and non-gen. models, these graph-related control RDMs were designed to capture more general structural properties of the graphs, without a specific distinction between the generalisable and non-generalisable feature dimension. The link distance, average bidirectional node distance and node type (hub, input, output) model RDMs were constructed as models of abstract graph coding, measuring abstract node relationships that transferred across block types (shape vs. texture) and environments (familiar vs. novel). The bidirectional one-step transition model was the opposite, designed to capture local node relationships specific to each block type-environment combination. We further included participant specific control RDMs to capture the average difference in accuracy between condition pairs and differences in the number of visual stimulus matches shown during the response and feedback screens. Control RDMs are visualised in Figure 3.

Paralleling the behavioural analyses, we first considered data from all trials (Fig. 6A). We began with a 2×2 repeated measures ANOVA on the RSA coefficient magnitudes, with factors of model (gen. vs. non. gen.) and ROI (EC vs. HPC). The analysis did not detect significant main effects of model (*F*(1,39)=1.32, *p*=0.257) or ROI (*F*(1,39)=1.95, *p*=0.171), and did not detect a significant interaction between model and ROI (*F*(1,39)=0.05, *p*=0.817). This indicates that the RSA model coefficients estimated using all trials in the novel environment did not reliably differ as function of generalisability or ROI. To test whether each RSA model itself was a significant predictor of each neural RDM, we conducted a series of permutation tests. These indicated that the gen. and non-gen model RDMs were both significant predictors of the neural data from EC (mean *β*_gen_=0.020, SD_gen_=0.040, permutation test: *p*=0.002; mean *β*_non-gen_=0.013, SD_non-gen_=0.028, permutation test: *p*=0.002, Fig. 6A). Results were the same for HPC (mean *β*_gen_=0.027, SD_gen_=0.043, permutation test: *p*<0.001; mean *β*_non-gen_=0.019, SD_non-gen_=0.030, permutation test: *p*<0.001). Together, these findings suggest that, on average across the full session, EC and HPC used similar neural codes to represent the cued stimulus feature across familiar and novel environments when sequences did and did not generalise.

**Figure 6.**
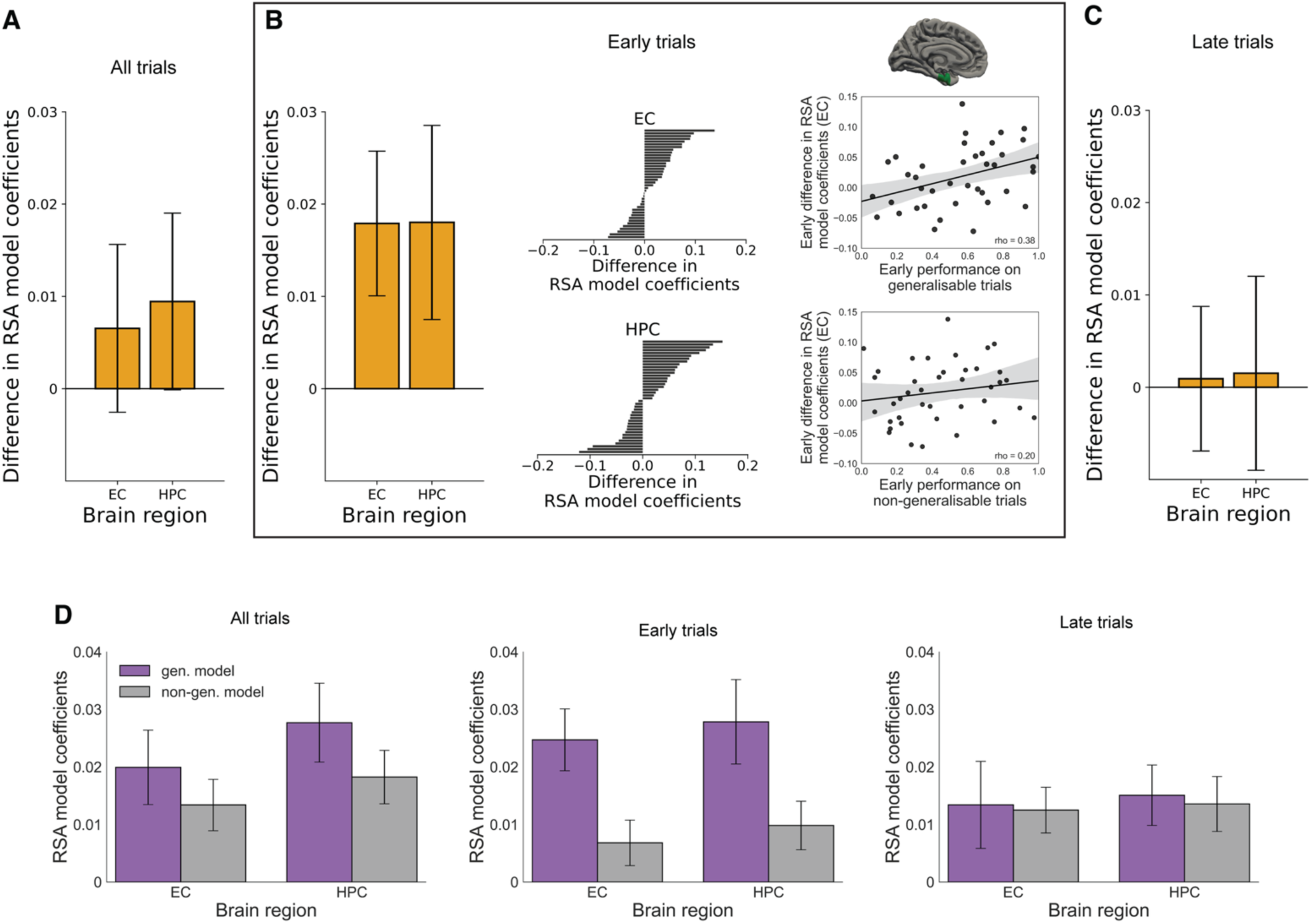
Neural results. **A:** The difference in RSA model coefficients between the gen. and non-gen. model RDMs for entorhinal cortex (EC) and hippocampus (HPC). A higher difference score indicates greater neural similarity across familiar and novel environments for the generalisable feature dimension. Bars show standard error of the mean. **B:** The same panel structure as A but showing coefficient differences for early trials in the novel environment (trials 1-80). Tornado plots show difference scores for individual participants. Scatterplots show correlations between the early neural effect in EC and performance in the novel environment. Performance is separated into early trials where the sequences from familiar environment either could or could not be generalised. Dots show individual participant values. Black lines indicate linear fits the data and gray lines indicate 95% confidence intervals of the fits. **D:** RSA coefficients for the gen. and non-gen. models during all trials (1-160), early trials in the novel environment (1-80) and late trials in the novel environment (81-160). Bars show standard error of the mean.

We next considered early exposure to the novel environment (the first half of the scanning session), where there was a prominent difference in behavioural performance (Fig. 5E). In this case, a 2×2 repeated measures ANOVA on coefficient magnitude, with factors of model (gen. vs. non-gen.) and ROI (EC vs. HPC) revealed a significant main effect of model RDM (*F*(1,39)=5.50, *p*=0.024), reflecting higher coefficient magnitudes for the gen. model RDM (M=0.03, SD=0.04) compared to the non-gen. model RDM (M=0.01, SD=0.03). The ANOVA did not detect a significant main effect of ROI (*F*(1,39)=0.60, *p*=0.445) or an interaction between model RDM and ROI (*F*(1,39)<0.01, *p*=0.991). Although the ANOVA detected a significant effect of model but not a significant interaction between model and ROI, we were motivated to proceed with permutation tests of the coefficient difference within each ROI, due to the previous theoretical proposal that EC factorises multi-dimensional events (Behrens et al., 2018; Whittington et al., 2022) and might therefore be attuned to the distinction between generalisable and non-generalisable dimensions. Permutation tests of the model RDM difference within each ROI indicted that coefficients for the gen. model fit to the EC data were significantly higher than coefficients for the non-gen. model (M_diff_=0.018, SD_diff_=0.049, permutation test: *p*=0.026, Fig. 6B). Coefficients were not significantly different between RSA models fit to the HPC data (M_diff_=0.018, SD_diff_=0.066, permutation test: *p*=0.104, Fig. 6B). We next tested whether each RSA model itself was a significant predictor of each neural RDM. This indicated that the gen. model RDM was a significant predictor of the neural data from EC (mean *β*_gen_=0.025, SD_gen_=0.034, permutation test: *p*<0.001, Fig. 6B) but the non-gen. model RDM did not reach significance (mean *β*_non-gen_=0.007, SD_non-gen_=0.025, permutation test: *p*=0.062, Fig. 6B). Equivalent tests indicated that the gen. and non-gen. models were both significant predictors of the neural data from HPC (mean *β*_gen_=0.028, SDgen=0.046, permutation test: *p*<0.001; mean *β*_non-gen_=0.010, SD_non-gen_=0.026, permutation test: *p*=0.022). Coefficients for all RSA models fit to the neural RDMs for EC and HPC during these analyses are shown in Fig. S1. An RSA analysis that used novel environment trials from the late phase (trial 81-160 of 160) rather than the early phase did not show significant differences between the gen. and non-gen. models (EC: M_diff_=0.001, SD_diff_=0.057, permutation test: *p*=0.925, HPC: M_diff_=0.002, SD_diff_=0.048, permutation test: *p*=0.864, Fig. 6C). The numerical reduction between the gen. and non-gen. coefficients from the early phase to the late phase did not reach significance (EC: M_diff_=-0.02, SD_diff_=0.07, permutation test: *p*=0.133, HPC: M_diff_=-0.02, SD_diff_=0.08, permutation test: *p*=0.182). Together, the collection of results in this section suggests that neural patterns in the EC distinguished early trials in the novel environment where sequences from the familiar environment could be generalised (i.e. the gen. and non-gen. model coefficients were significantly different from each other in the EC). However, it was also the case that this neural signalling profile was not significantly higher in EC compared to HPC (i.e. the difference between the gen. and non-gen. coefficients was not significantly different between the ROIs).

The results so far provide evidence for a neural distinction between trials involving the generalisable and non-generalisable feature dimensions during early exposure to the novel environment. These results were based on model RDMs that quantified the neural similarity between stimuli at the starting node across trials. As a next step, we tested whether the distinction between the gen. and non-gen. conditions extended to the next stimulus in the sequence. To do so, we repeated the analysis but added three model RDMs to account for cross-environment forward transitions (Fig. 4). The main idea was that when an animal cue was shown in the novel environment, the stimulus at the next node in the graph would be activated. This activation would increase the neural similarity between the current condition in the novel environment and the condition in the familiar environment that began with that activated feature. For example, seeing an elephant cue in the novel environment during a shape block (the circle node in Fig. 1), would lead to the activation of the triangle as the next node, increasing the similarity with lemur trials in the familiar environment where the triangle was cued. One model RDM captured such one-step forward transitions for the generalisable blocks of the novel environment. Another captured one-step forward transitions for the non-generalisable blocks. A final model RDM captured one-step forward transitions in the non-generalisable blocks based on the irrelevant graph structure from the familiar environment. A 3×2 repeated measures ANOVA on coefficient magnitude, with factors of model (gen. forward vs. non-gen. forward vs. non-gen. irrel. forward) and ROI (EC. vs. HPC) did not detect significant main effects of model (*F*(2,78)=0.017, *p*=0.890), ROI (*F*(1,39)=3.78, *p*=0.059) or a significant interaction between model and ROI and block (*F*(2,78)=2.24, *p*=0.113). Coefficients for all RSA models fit to the neural RDMs for EC and HPC during these analyses are shown in Fig. S2.

We explored two possibilities for the absence of a significant difference between forward transition models. One possibility is that a distinction between the generalisable and non-generalisable feature dimension did extend to the next node in the sequence, but later in the trial. To test this, we re-ran the analysis using data locked to the response phase, when one-step transitions needed to be applied (+15 to +20s from trial onset, including a shift of 3TRs to account for hemodynamic lag). The corresponding 3×2 repeated measures ANOVA on coefficient magnitude, with factors of model (gen. forward vs. non-gen. forward vs. non-gen. irrel. forward) and ROI (EC. vs. HPC) did not detect significant main effects of model (*F*(2,78)=0.11, *p*=0.890), ROI (*F*(1,39)=2.36, *p*=0.133) or a significant interaction between ROI and block (*F*(2,78)=1.04, *p*=0.358). A second possibility is that differences between forward transition models were transient. Averaging over the full decision phase might have masked these differences. To test this idea, we used cluster-based permutation testing, a method capable of detecting transient differences between time courses (Maris & Oostenveld, 2007). No significant differences between forward transition models were detected across the planning and response phases (testing window: +3 to +20s from trial onset, EC gen. forward model vs. EC non-gen. forward model: candidate cluster at 5s, *p*=0.277; EC gen. forward model vs. EC non-gen. irrel. forward model: candidate clusters at 5s, *p*=0.211, and 11.25s, *p*=0.128; HPC gen. forward model vs. HPC non-gen. forward model: no candidate clusters; HPC gen. forward model vs. HPC non-gen. irrel. forward model: candidate cluster at 12.5s, *p*=0.308). Summarising the results above, we did not detect evidence that the neural differentiation between generalisable and non-generalisable conditions (Fig. 6B) extended to the second node in each sequence during early trials in the novel environment.

We next sought to explore whether the neural differentiation between the gen. and non-gen. conditions during early trials (Fig. 6B) was related to activity in other brain regions that have been associated with graph-like representations or the generalisation of latent task structure. These included the medial orbitofrontal cortex (Schuck et al., 2016; Park et al., 2020), the rostral anterior cingulate cortex (Baram et al., 2024), as well as the dorsolateral prefrontal cortex, inferior parietal cortex and precuneus (Vaidya et al., 2021). To do so, we re-ran our analysis pipeline to estimate gen. and non-gen. coefficients based on early trials in the novel environment, for each exploratory region. We then conducted a 2×7 repeated measures ANOVA on coefficient magnitude, with factors of model (gen. vs non-gen.) and ROI (the five exploratory regions, plus EC and HPC). The ANOVA indicated a significant main effect of ROI (*F*(6,234)=2.31, *p*=0.035) but no significant main effect of model (*F*(1,39)=1.93, *p*=0.173) or interaction between model and ROI (*F*(6,234)=1.90, *p*=0.081). The absence of a significant interaction between model and ROI indicates that the early generalisation signal in EC (Fig. 6B, Fig. S3) was not significantly larger than in the other brain regions we examined. Coefficients for the full set of ROIs are presented in Fig. S4 and S5.

To conclude our investigation, we examined functional connections between neural signalling and behaviour. Given the neural results above, the correlation tests that follow were all conducted using data from early trials in the novel environment. We first focused on whether the difference in neural signalling between gen. and non-gen. conditions during early trials (Fig. 6B) tracked the corresponding difference in behavioural performance (Fig. 5E). We did not detect significant Spearman correlations between these variables for the EC (*ρ*(38)=0.29, *p*=0.133) or the HPC (*ρ*(38)=0.05, *p*=0.749). We next considered performance on the generalisable and non-generalisable trials separately. In this case, stronger neural signalling in EC was associated with significantly higher accuracy on generalisable trials, where prior sequence knowledge could be reused (*ρ*(38)=0.38, *p*=0.029, Fig. 6B). The equivalent neuro-behavioural association was not significant for non-generalisable trials, which required new sequential transitions (*ρ*(38)=0.20, *p*=0.440, Fig. 6B). A Williams test for overlapping correlations indicated that the EC neuro-behavioural correlation for generalisable trials was not significantly stronger than the correlation for non-generalisable trials (*t*(37)=1.62, *p*=0.114). For HPC, no significant associations were detected between neural signalling (Fig. 6B) and performance on generalisable trials (*ρ*(38)=-0.05, *p*=0.762) or non-generalisable trials (*ρ*(38)=-0.10, *p*=0.549). The HPC neuro-behavioural correlation for generalisable trials was found to be significantly weaker than the equivalent neuro-behavioural correlation for EC (Williams test: *t*(37)=-2.61, *p*=0.013). Neuro-behavioural correlations for the exploratory ROIs are presented in Fig. S6. The most notable result from those analyses was that differential neural signalling in the precuneus also showed a significant correlation with performance in the generalisable blocks (*ρ*(38)=0.48, *p*=0.003, Fig. S6), despite the absence of a group-level difference between the gen. and non-gen. model coefficients (Fig. S3-S5). Remaining neuro-behavioural correlations were non-significant (*ρ*(38)<0.27, all *p*-values>0.19, Fig. S6). These findings suggest that neural patterns in the EC and precuneus, distinguishing the generalisable and non-generalisable feature dimensions, were associated with improved performance when prior sequences could be reused.

## Discussion

A central problem we face when attempting to generalise our experience to new situations, is determining which aspects of a new situation are similar to what we have experienced before, which are different, and which similarities matter for our current goals. This allows us to generalise our knowledge to the aspects that are similar and relevant for goal pursuit, but avoid over-generalising and set to work learning anew any critical aspects that differ. We investigated how this selective generalisation problem is overcome. Participants distinguished information shared across environments, with higher performance on blocks where previously learned sequences could be reused. This behavioural distinction was prominent during early exposure to the novel environment. Throughout this early phase, neural patterns in EC showed a corresponding differentiation between the shared and unshared feature dimensions. Neural patterns in EC representing starting locations in the familiar and novel graphs were significantly more similar on trials where sequences could be generalised from prior experience. This differential signalling was associated better performance on generalisable task blocks. These results point towards a possible neural mechanism for choice in novel environments, in which EC signals dimensions of past experience for reuse.

Previous theoretical work has proposed that EC has a role in factorising multi-dimensional events (Behrens et al., 2018; Whittington et al., 2022), a function which could help to distinguish generalisable and non-generalisable dimensions across familiar and novel environments. The present results appear to provide some empirical support for this notion. The theoretical work above also proposes that factorised representations in EC are distinct from representations in HPC, a region more often associated with conjunctive coding (see Komorowski et al., 2009; Rudy & O’Reilly, 2001; Zheng et al., 2024). We did not observe supporting evidence for this latter claim in our data, as a significant difference was not detected in the neural signalling profile between HPC and EC. One relevant question to consider is whether our experimental design inadvertently favoured a factorised solution by asking participants to separately report shape and texture sequences on different trials. In principle, participants could represent sequence information in either a conjunctive or factorised format in the familiar environment, since reactivating one or both features at each node during planning or response execution would not actively interfere with behavioural accuracy. The conjunctive solution in this case might be considered more efficient (one needs just 5 memory slots to represent the graph structure) but less flexible (one will experience interference if features are later recombined as was the case in the novel environment). The factorised solution has the opposite pros and cons. It might be considered less efficient (one needs 10 memory slots to represent all of the node information) but this information can be then used more flexibly with minimal interference, if features are later recombined. With these considerations in mind, it is not a given that a factorised solution would be the default representation when people need to report feature dimensions separately (provided there is no interference). Nonetheless, the converse is not true either. We cannot rule out the possibility that having participants report different feature dimensions in different trials did not contribute to a corresponding separation in how the feature information was represented neurally.

Previous neuroimaging studies have demonstrated that EC clusters information based on structural relationships when participants have extensive pretraining (Baram et al., 2021; Mark et al., 2024; Shpektor et al., 2024) or explicit instruction about these relationships (Glitz et al., 2022). In these cases, neural patterns in EC are more similar in response to stimuli that have the same abstract meaning (Shpektor et al., 2024) or correlated reward contingencies (Baram et al., 2021; Glitz et al., 2022). These results provide insight into the neural mechanisms that underpin certain kinds of generalisation behaviour. For example, such neural clustering could allow observed changes in the reward or transition probabilities of one stimulus to be applied to related stimuli. Nonetheless, there are important forms of generalisation that cannot be studied with extensive pretraining or explicit instruction. For example, when a person moves to a new country, they are unlikely to have full knowledge about their new environment on arrival. Instead, they must rapidly discern which aspects of their earlier experiences can be applied to make effective decisions in their new context. We attempted to create an experimental version of this generalisation problem, in which people have extensive experience with a familiar environment but do not know the structure of a novel environment in advance. Under these conditions, EC appears to play an early role in distinguishing the dimensions of past experience that can be generalised, through relative changes in representational similarity that correlate with improved choice performance.

The current experiment also differed from previous neuroimaging studies on generalisation in that participants had to transfer one feature dimension across contexts, but ignore another. This is reminiscent of cognitive control studies investigating how task-relevant stimulus features are processed while irrelevant features are supressed (e.g. Johnson et al., 2023; Richter et al., 2025; Ritz & Shenhav, 2024). While some findings from this domain suggest a persistent influence of task-irrelevant features on behaviour (Moneta et al., 2023; Tankelevitch et al., 2020), participants in the current experiment quickly learned to refrain from reusing transitions that were not applicable in the novel environment. To further understand generalisation to multidimensional environments, it could therefore be valuable for future research to consider intersections between factorised representations and putative cognitive control mechanisms for goal-directed processing of relevant feature dimensions, such as attentional gating (Johnson et al., 2023), proactive control (Richter et al., 2025) or neural geometry (Badre, 2024).

Another direction for future research based on the current results is to understand how early feedback in novel environments contributes to the formation of generalisation signals in EC. The present work does not address the nature of this connection because the trial-by-trial resolution of the neural analysis pipeline is limited by various factors. One is that all conditions are represented in the neural RDM. This means that the resulting RSA coefficients reflect information from multiple trials and do not provide neural coding estimates for individual trials. A second factor is that the pipeline relies on averaging trials for each condition to reduce noise and compute robust estimates of cross-environment coding patterns for the generalisable and non-generalisable conditions. Our approach therefore lacks a more fine-grained trial-by-trial resolution that would be ideal to relate neural signals involved in feedback processing (e.g. prediction errors) and evolving generalisation signals across learning. One strategy for addressing this in the future could be to combine reinforcement learning models with emerging methods for single-trial RSA analysis (e.g. Huang et al., 2025), to estimate trialwise prediction errors in a novel environment and test whether these relate to trialwise changes in cross-environment neural similarity. In addition to gaining better mechanistic insight into the specific relationships between feedback in novel environments and the development generalisation signals in EC, the neural effects observed in the present dataset were small. It would therefore be valuable in general to replicate these EC signals in a future experiment with a simpler design, to confirm these changes during transfer to novel environments.

The hypothesis tested in this work is that EC contributes to selective generalisation by signalling the difference between generalisable and non-generalisable dimensions in novel environments. A more extreme hypothesis is that this function is exclusive to EC. The present results provide some initial evidence for the primary hypothesis, but do not provide evidence for the more extreme hypothesis. Although EC was the only region we examined that showed both a significant neural distinction between generalisable and non-generalisable dimensions (Fig. S3) and a significant correlation with generalisation performance (Fig. 6B, Fig. S6), its neural effects were not significantly stronger than other brain regions associated with cognitive maps and generalisation. Therefore, we cannot conclude that neural signalling of generalisable dimension was specific or exclusive to EC. The observation that generalisation signals in EC were not statistically distinct from the other brain areas we examined raises interesting questions. For instance, it remains open as to whether these other brain regions – especially those close to the threshold for significance such as mOFC (Fig. S3) or in close anatomical proximity such as HPC - carry similar generalisation signals to EC. If this is the case, an interesting question to resolve would be whether this due to local factorisation processes in each region or because these regions receive input from EC about which dimensions generalise.

Another intriguing finding about the ROIs we examined beyond EC was that the precuneus showed the lowest group-level distinction between the generalisable and non-generalisable dimensions (Fig. S3). Yet, individual differences in precuneus signalling were significantly correlated with performance on the generalisation blocks (Fig. S6). One speculation is this may reflect distinct generalisation strategies that recruit the precuneus to different extents and confer different degrees of generalisation success in multi-dimensional environments. Exploring which generalisation strategies precuneus recruitment relates to (if any) is an intriguing area for future investigation.

The present results only support limited conclusions about the role of EC in planning, as a specific use case of selective generalisation. We did not detect reliable evidence that the distinction between the generalisable and non-generalisable feature dimensions extended past the starting location, to the next node in the sequence. One possibility could be that the visual presentation of the starting animal induced relatively strong neural responses across participants, due to well-learned associations between each animal cue and the relevant stimulus feature at that node in the graph. In contrast, internal planning operations from the starting point to subsequent nodes may have been weaker and more variable across participants, making it harder to detect effects at the group level. Although an effect of generalisability was not detected past the starting location, the results showed a significant correlation between EC generalisation signalling and planning performance on the early generalisable blocks, which suggests a connection between EC signalling and planning. While these differing results make it hard to draw firm conclusions about EC’s role in planning, we speculate that EC’s potential role in selective generalisation could be a more general function that is not limited to planning scenarios. This possibility could be tested in the future, with tasks that require selective generalisation but not sequential planning.

Overall, EC exhibited neural signalling consistent with selective generalisation and a correlation between these signals and generalisation performance. These findings do not imply that EC only encodes the distinction between generalisable and non-generalisable dimensions or that EC is the only region that contributes to selective generalisation. The present results suggest that during early exposure to novel environments, EC may signal dimensions of past experience that can be generalised. Ultimately, such a signal would be expected to aid choice performance in novel settings, allowing us to discern between reusing past experience and learning new environmental structures.

## Acknowledgements

This research was supported by a New Zealand Neurological Foundation Fellowship and an Alexander von Humboldt Fellowship awarded to Sam Hall-McMaster, an Independent Max Planck Research Group grant and a Starting Grant from the European Union (ERC-2019-StG REPLAY-852669) awarded to Nicolas W. Schuck, and a Multi-disciplinary University Research Initiative (MURI) award by the Army Research Office (W911NF-21-1-0328 and W911NF-23-1-0277) to Samuel J. Gershman. Peter Dayan was funded by the Max Planck Society and the Humboldt Foundation.

## References

Abraham, A., Pedregosa, F., Eickenberg, M., Gervais, P., Mueller, A., Kossaifi, J., … & Varoquaux, G. (2014). Machine learning for neuroimaging with scikit-learn. Frontiers in Neuroinformatics, 8, 14. 10.3389/fninf.2014.00014

Avants, B. B., Epstein, C. L., Grossman, M., & Gee, J. C. (2008). Symmetric diffeomorphic image registration with cross-correlation: Evaluating automated labeling of elderly and neurodegenerative brain. Medical Image Analysis, 12(1), 26–41.

Badre, D. (2024). Cognitive control. Annual Review of Psychology, 76.

Baram, A. B., Muller, T. H., Nili, H., Garvert, M. M., & Behrens, T. E. J. (2021). Entorhinal and ventromedial prefrontal cortices abstract and generalize the structure of reinforcement learning problems. Neuron, 109(4), 713–723.

Baram, A., Nili, H., Barreiros, I., Samborska, V., Behrens, T. E., & Garvert, M. M. (2024). An abstract relational map emerges in the human medial prefrontal cortex with consolidation. bioRxiv, 2024-10.

Bellmund, J. L., Deuker, L., Montijn, N. D., & Doeller, C. F. (2022). Mnemonic construction and representation of temporal structure in the hippocampal formation. Nature Communications, 13(1), 3395.

Benjamini, Y., & Hochberg, Y. (1995). Controlling the false discovery rate: A practical and powerful approach to multiple testing. Journal of the Royal Statistical Society: Series B (Methodological), 57(1), 289–300.

Behrens, T. E., Muller, T. H., Whittington, J. C., Mark, S., Baram, A. B., Stachenfeld, K. L., & Kurth-Nelson, Z. (2018). What is a cognitive map? Organizing knowledge for flexible behavior. Neuron, 100(2), 490–509.

Behzadi, Y., Restom, K., Liau, J., & Liu, T. T. (2007). A component based noise correction method (CompCor) for BOLD and perfusion based fMRI. Neuroimage, 37(1), 90–101.

Bhui, R. (2018). Case-based decision neuroscience: Economic judgment by similarity. In Goal-directed decision making (pp. 67–103). Academic Press.

Brunec, I. K., & Momennejad, I. (2022). Predictive representations in hippocampal and prefrontal hierarchies. Journal of Neuroscience, 42(2), 299–312.

Collins, A. G., & Frank, M. J. (2013). Cognitive control over learning: creating, clustering, and generalizing task-set structure. Psychological Review, 120(1), 190.

Cox, R. W., & Hyde, J. S. (1997). Software tools for analysis and visualization of fMRI data. NMR in Biomedicine, 10(4-5), 171–178. 10.1002/(SICI)1099-1492(199706/08)10:4/5<171::AID-NBM453>3.0.CO;2-L

Dale, A. M., Fischl, B., & Sereno, M. I. (1999). Cortical surface-based analysis: I. Segmentation and surface reconstruction. Neuroimage, 9(2), 179–194. 10.1006/nimg.1998.0395

Esteban O, Birman D, Schaer M, Koyejo OO, Poldrack RA, Gorgolewski KJ; MRIQC: Advancing the Automatic Prediction of Image Quality in MRI from Unseen Sites; Plos One 12(9):e0184661; doi:10.1371/journal.pone.0184661.

Esteban, Oscar, Christopher Markiewicz, Ross W Blair, Craig Moodie, Ayse Ilkay Isik, Asier Erramuzpe Aliaga, James Kent, et al. 2018. “fMRIPrep: A Robust Preprocessing Pipeline for Functional MRI.” Nature Methods. 10.1038/s41592-018-0235-4.

Esteban, O., Markiewicz, C. J., Goncalves, M., Provins, C., Salo, T., Kent, J. D., DuPre, E., Ciric, R., Pinsard, B., Blair, R. W., Poldrack, R. A., & Gorgolewski, K. J. (2024). fMRIPrep: A robust preprocessing pipeline for functional MRI (23.2.1). Zenodo. 10.5281/zenodo.10790684

Evans, A. C., Janke, A. L., Collins, D. L., & Baillet, S. (2012). Brain templates and atlases. Neuroimage, 62(2), 911–922.

Flesch, T., Juechems, K., Dumbalska, T., Saxe, A., & Summerfield, C. (2022). Orthogonal representations for robust context-dependent task performance in brains and neural networks. Neuron, 110(7), 1258–1270.

Fonov, V. S., Evans, A. C., McKinstry, R. C., Almli, C. R., & Collins, D. L. (2009). Unbiased nonlinear average age-appropriate brain templates from birth to adulthood. Neuroimage, 47, S102. 10.1016/S1053-8119(09)70884-5

Garvert, M. M., Dolan, R. J., & Behrens, T. E. (2017). A map of abstract relational knowledge in the human hippocampal–entorhinal cortex. elife, 6, e17086.

Glitz, L., Juechems, K., Summerfield, C., & Garrett, N. (2022). Model sharing in the human medial temporal lobe. Journal of Neuroscience, 42(27), 5410–5426.

Gorgolewski, K., Burns, C. D., Madison, C., Clark, D., Halchenko, Y. O., Waskom, M. L., & Ghosh, S. S. (2011). Nipype: a flexible, lightweight and extensible neuroimaging data processing framework in python. Frontiers in Neuroinformatics, 5, 13. 10.3389/fninf.2011.00013

Gorgolewski, Krzysztof J., Oscar Esteban, Christopher J. Markiewicz, Erik Ziegler, David Gage Ellis, Michael Philipp Notter, Dorota Jarecka, et al. 2018. Nipype. *Zenodo*. 10.5281/zenodo.596855

Greve, D. N., & Fischl, B. (2009). Accurate and robust brain image alignment using boundary-based registration. Neuroimage, 48(1), 63–72. 10.1016/j.neuroimage.2009.06.060.

Gustafson, N. J., & Daw, N. D. (2011). Grid cells, place cells, and geodesic generalization for spatial reinforcement learning. Plos Computational Biology, 7(10), e1002235.

Hall-McMaster, S., Stokes, M. G., & Myers, N. E. (2022). Integrating Reward Information for Prospective Behavior. Journal of Neuroscience, 42(9), 1804–1819.

Huang, S., Howard, C. M., Bogdan, P. C., Morales-Torres, R., Slayton, M., Cabeza, R., & Davis, S. W. (2025). Trial-level representational similarity analysis. BioRxiv, 2025-03.

Jenkinson, M., Beckmann, C. F., Behrens, T. E., Woolrich, M. W., & Smith, S. M. (2012). Fsl. Neuroimage, 62(2), 782–790.

Jenkinson, M., Bannister, P., Brady, M., & Smith, S. (2002). Improved optimization for the robust and accurate linear registration and motion correction of brain images. Neuroimage, 17(2), 825–841.

Johnson, E. L., Lin, J. J., King-Stephens, D., Weber, P. B., Laxer, K. D., Saez, I., … & Badre, D. (2023). A rapid theta network mechanism for flexible information encoding. Nature Communications, 14(1), 2872.

Klein, A., Ghosh, S. S., Bao, F. S., Giard, J., Häme, Y., Stavsky, E., … & Keshavan, A. (2017). Mindboggling morphometry of human brains. Plos Computational Biology, 13(2), e1005350.

Krüger, G., & Glover, G. H. (2001). Physiological noise in oxygenation-sensitive magnetic resonance imaging. Magnetic Resonance in Medicine: An Official Journal of the International Society for Magnetic Resonance in Medicine, 46(4), 631–637.

Lanczos, C. (1964). Evaluation of noisy data. Journal of the Society for Industrial and Applied Mathematics, Series B: Numerical Analysis, 1(1), 76–85.

Lindsey, J. W., & Issa, E. B. (2024). Factorized visual representations in the primate visual system and deep neural networks. Elife, 13, RP91685.

Liu, Y., Dolan, R. J., Kurth-Nelson, Z., & Behrens, T. E. (2019). Human replay spontaneously reorganizes experience. Cell, 178(3), 640–652.

Luyckx, F., Nili, H., Spitzer, B., & Summerfield, C. (2019). Neural structure mapping in human probabilistic reward learning. elife, 8, e42816.

Manns, J. R., & Eichenbaum, H. (2006). Evolution of declarative memory. Hippocampus, 16(9), 795–808.

Maris, E., & Oostenveld, R. (2007). Nonparametric statistical testing of EEG-and MEG-data. Journal of Neuroscience Methods, 164(1), 177–190.

Mark, S., Schwartenbeck, P., Hahamy, A., Samborska, V., Baram, A. B., & Behrens, T. E. J. (2023). Flexible neural representations of abstract structural knowledge in the human Entorhinal Cortex. bioRxiv, 2023-08.

Moneta, N., Garvert, M. M., Heekeren, H. R., & Schuck, N. W. (2023). Task state representations in vmPFC mediate relevant and irrelevant value signals and their behavioral influence. Nature Communications, 14(1), 3156.

Myers, N. E., Rohenkohl, G., Wyart, V., Woolrich, M. W., Nobre, A. C., & Stokes, M. G. (2015). Testing sensory evidence against mnemonic templates. elife, 4, e09000.

O’Reilly, R. C., & Rudy, J. W. (2001). Conjunctive representations in learning and memory: principles of cortical and hippocampal function. Psychological Review, 108(2), 311.

Park, S. A., Miller, D. S., Nili, H., Ranganath, C., & Boorman, E. D. (2020). Map making: constructing, combining, and inferring on abstract cognitive maps. Neuron, 107(6), 1226–1238.

Power, J. D., Barnes, K. A., Snyder, A. Z., Schlaggar, B. L., & Petersen, S. E. (2012). Spurious but systematic correlations in functional connectivity MRI networks arise from subject motion. Neuroimage, 59(3), 2142–2154.

Komorowski, R. W., Manns, J. R., & Eichenbaum, H. (2009). Robust conjunctive item–place coding by hippocampal neurons parallels learning what happens where. Journal of Neuroscience, 29(31), 9918–9929.

Koolschijn, R. S., Emir, U. E., Pantelides, A. C., Nili, H., Behrens, T. E., & Barron, H. C. (2019). The hippocampus and neocortical inhibitory engrams protect against memory interference. Neuron, 101(3), 528–541.

Kriegeskorte, N., Mur, M., & Bandettini, P. A. (2008). Representational similarity analysis-connecting the branches of systems neuroscience. Frontiers in Systems Neuroscience, 2, 249.

Reuter, M., Rosas, H. D., & Fischl, B. (2010). Highly accurate inverse consistent registration: a robust approach. Neuroimage, 53(4), 1181–1196.

Richter, D., van Moorselaar, D., & Theeuwes, J. (2025). Proactive distractor suppression in early visual cortex. Elife, 13, RP101733.

Ritz, H., & Shenhav, A. (2024). Orthogonal neural encoding of targets and distractors supports multivariate cognitive control. Nature Human Behaviour, 8(5), 945–961.

Saad, Z. S., Reynolds, R. C., Jo, H. J., Gotts, S. J., Chen, G., Martin, A., & Cox, R. W. (2013). Correcting brain-wide correlation differences in resting-state FMRI. Brain Connectivity, 3(4), 339–352.

Satterthwaite, T. D., Elliott, M. A., Gerraty, R. T., Ruparel, K., Loughead, J., Calkins, M. E., … & Wolf, D. H. (2013). An improved framework for confound regression and filtering for control of motion artifact in the preprocessing of resting-state functional connectivity data. Neuroimage, 64, 240–256. 10.1016/j.neuroimage.2012.08.052

Schapiro, A. C., Kustner, L. V., & Turk-Browne, N. B. (2012). Shaping of object representations in the human medial temporal lobe based on temporal regularities. Current Biology, 22(17), 1622–1627.

Schapiro, A. C., Turk-Browne, N. B., Norman, K. A., & Botvinick, M. M. (2016). Statistical learning of temporal community structure in the hippocampus. Hippocampus, 26(1), 3–8.

Schuck, N. W., Cai, M. B., Wilson, R. C., & Niv, Y. (2016). Human orbitofrontal cortex represents a cognitive map of state space. Neuron, 91(6), 1402–1412.

Schütt, H. H., Kipnis, A. D., Diedrichsen, J., & Kriegeskorte, N. (2023). Statistical inference on representational geometries. Elife, 12, e82566.

Sharkey, N. E., & Sharkey, A. J. (1993). Adaptive generalisation. Artificial Intelligence Review, 7(5), 313–328.

Shepard, R. N. (1987). Toward a universal law of generalization for psychological science. Science, 237(4820), 1317–1323.

Shpektor, A., Bakermans, J. J., Baram, A. B., Sarnthein, J., Ledergerber, D., Imbach, L., … & Behrens, T. E. (2024). A hierarchical coordinate system for sequence memory in human entorhinal cortex. bioRxiv, 2024-10.

Tankelevitch, L., Spaak, E., Rushworth, M. F., & Stokes, M. G. (2020). Previously reward-associated stimuli capture spatial attention in the absence of changes in the corresponding sensory representations as measured with MEG. Journal of Neuroscience, 40(26), 5033–5050.

Tustison, N. J., Avants, B. B., Cook, P. A., Zheng, Y., Egan, A., Yushkevich, P. A., & Gee, J. C. (2010). N4ITK: improved N3 bias correction. IEEE Transactions on Medical Imaging, 29(6), 1310–1320. 10.1109/TMI.2010.2046908.

Weiskopf, N., Hutton, C., Josephs, O., & Deichmann, R. (2006). Optimal EPI parameters for reduction of susceptibility-induced BOLD sensitivity losses: a whole-brain analysis at 3 T and 1.5 T. Neuroimage, 33(2), 493–504.

Xia, L., & Collins, A. G. (2021). Temporal and state abstractions for efficient learning, transfer, and composition in humans. Psychological Review, 128(4), 643.

Vaidya, A. R., Jones, H. M., Castillo, J., & Badre, D. (2021). Neural representation of abstract task structure during generalization. ELife, 10, e63226.

Walther, A., Nili, H., Ejaz, N., Alink, A., Kriegeskorte, N., & Diedrichsen, J. (2016). Reliability of dissimilarity measures for multi-voxel pattern analysis. Neuroimage, 137, 188–200.

Wittkuhn, L., & Schuck, N. W. (2021). Dynamics of fMRI patterns reflect sub-second activation sequences and reveal replay in human visual cortex. Nature Communications, 12(1), 1795.

Whittington, J. C., McCaffary, D., Bakermans, J. J., & Behrens, T. E. (2022). How to build a cognitive map. Nature neuroscience, 25(10), 1257–1272.

Whittington, J. C., Muller, T. H., Mark, S., Chen, G., Barry, C., Burgess, N., & Behrens, T. E. (2020). The Tolman-Eichenbaum machine: unifying space and relational memory through generalization in the hippocampal formation. Cell, 183(5), 1249–1263.

Zhang, Y., Brady, M., & Smith, S. (2001). Segmentation of brain MR images through a hidden Markov random field model and the expectation-maximization algorithm. IEEE Transactions on Medical Imaging, 20(1), 45–57. 10.1109/42.906424.

Zheng, X. Y., Hebart, M. N., Grill, F., Dolan, R. J., Doeller, C. F., Cools, R., & Garvert, M. M. (2024). Parallel cognitive maps for multiple knowledge structures in the hippocampal formation. Cerebral Cortex, 34(2), bhad485.

