## Supplement for "Entorhinal cortex signals dimensions of past experience that can be generalised in a novel environment"

**Supplementary Information**

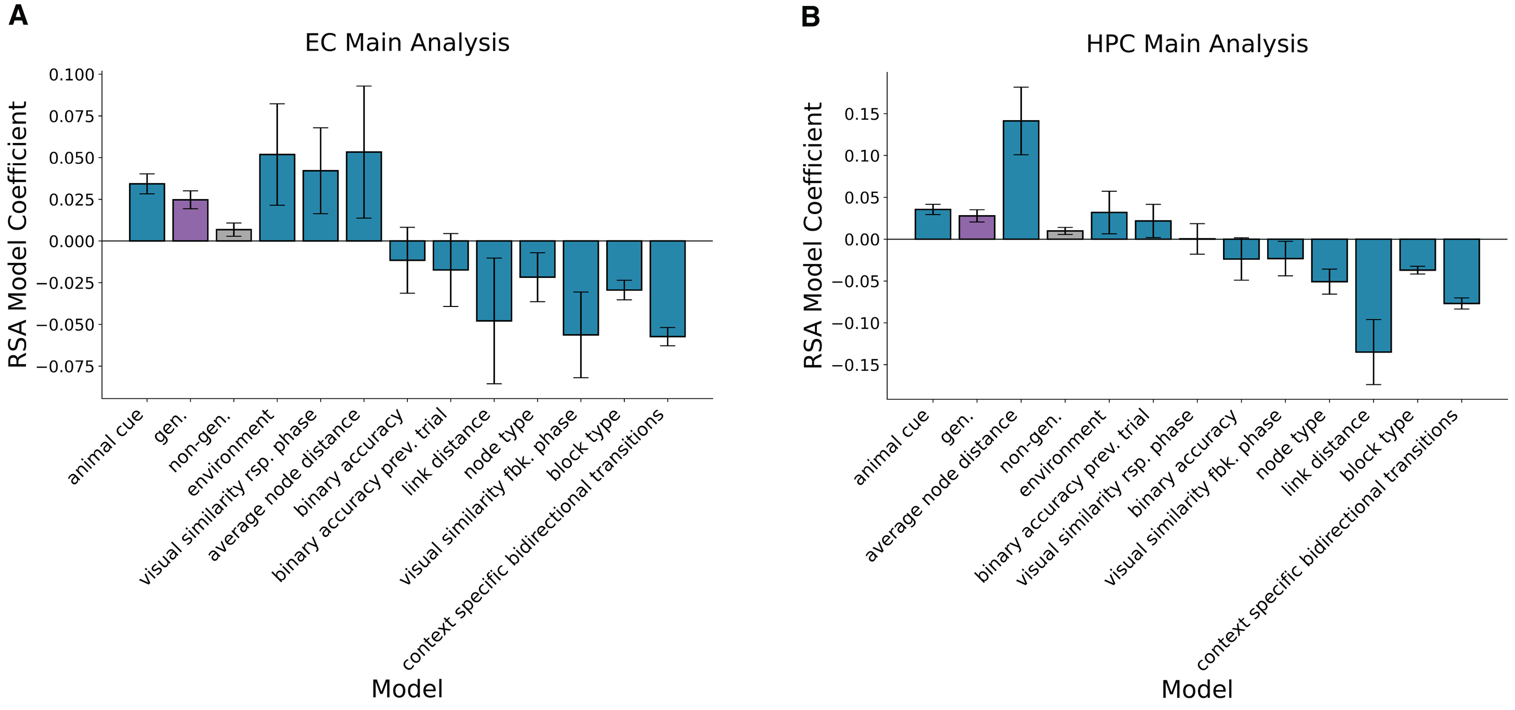

*Figure S1.* Coefficients for all representational similarity analysis (RSA) models included during the analysis of early trials in the novel environment. Model coefficients are based on an RSA that compared neural patterns during early trials in the novel environment (1-80) to neural patterns in the familiar environment (trials 1-160). Bars show the mean coefficient value and error bars show standard error of the mean. Bars are ordered left to right based on Cohen’s *d* effect sizes (mean coefficient value divided by standard deviation). The models include the gen. and non-gen. models shown in Fig. 2 of the main text and the control models shown in Fig. 3. **A:** Coefficients based on data from entorhinal cortex (EC). **B:** Coefficients based on data from hippocampus (HPC).

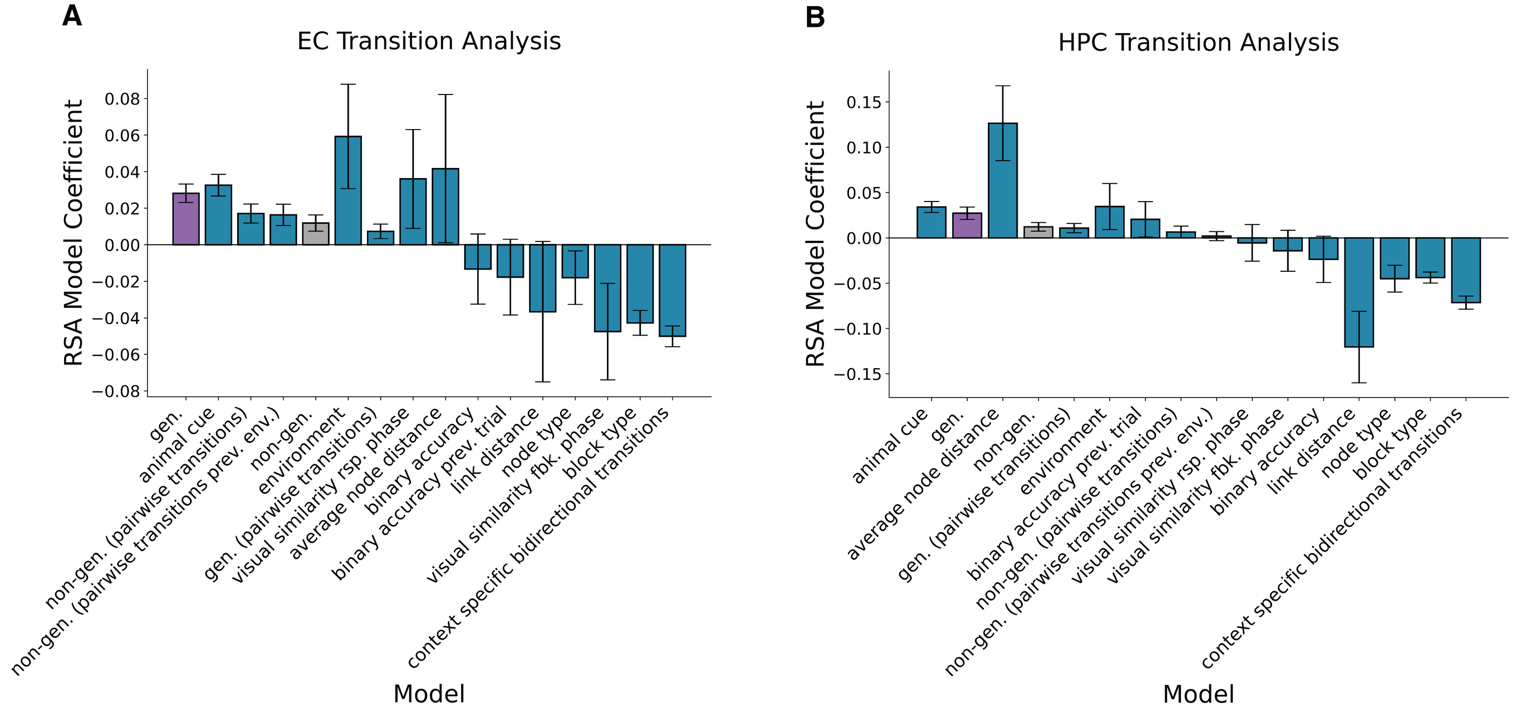

*Figure S2.* Coefficients for all representational similarity analysis (RSA) models included during forward transition analyses. Model coefficients are based on an RSA that compared neural patterns during early trials in the novel environment (1-80) to neural patterns in the familiar environment (trials 1-160). The RSA used neural data from the decision phase. Bars show the mean coefficient value and error bars show standard error of the mean. Bars are ordered left to right based on Cohen’s *d* effect sizes (mean coefficient value divided by standard deviation). The models include the core gen. and non-gen. models shown in Fig. 2 of the main text, the control models in Fig. 3 and the forward transition in Fig. 4. **A:** Coefficients based on data from entorhinal cortex (EC). **B:** Coefficients based on data from hippocampus (HPC).

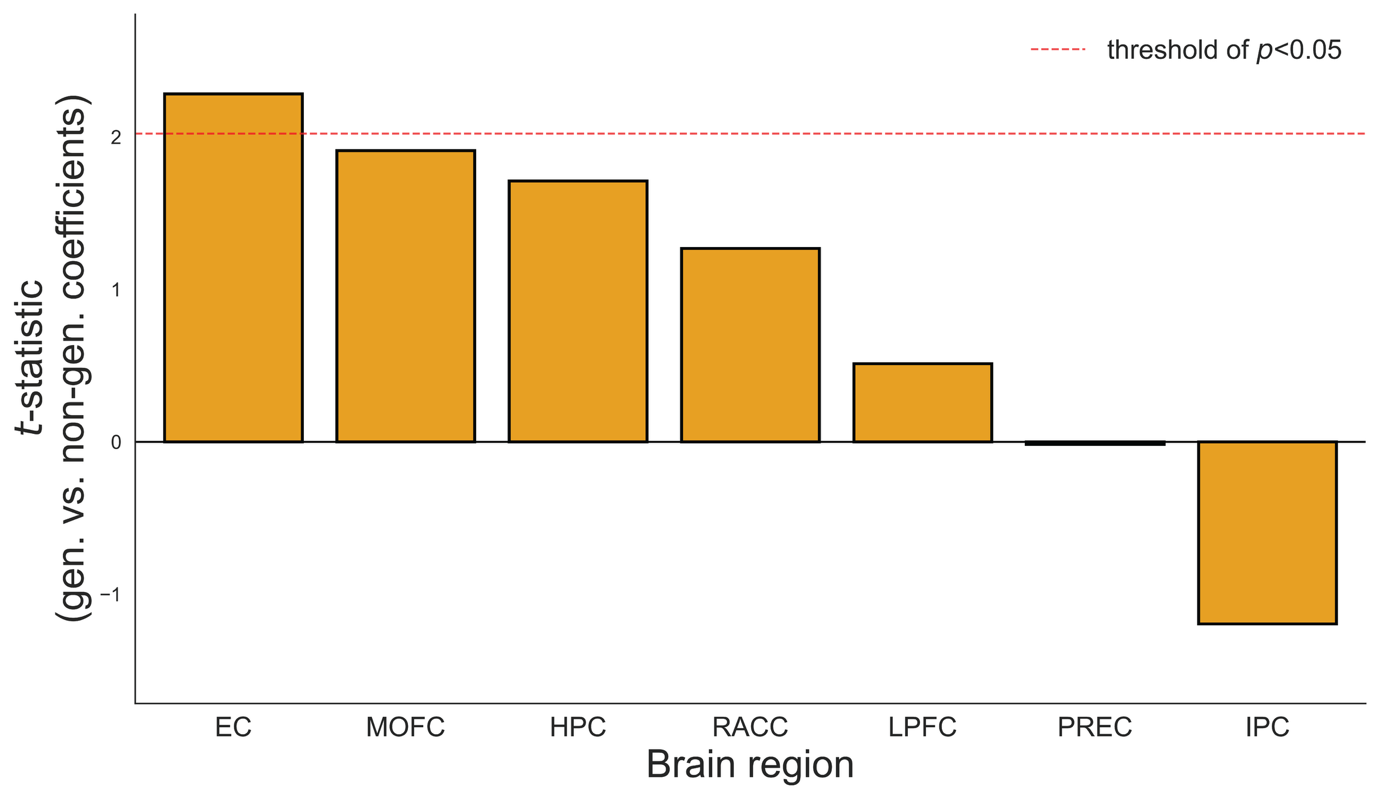

*Figure S3. t*-statistics for the difference between the gen. and non-gen. model coefficients, for all regions of interest. Model coefficients are based on representational similarity analyses that compare neural patterns during early trials in the novel environment (1-80) to neural patterns in the familiar environment (trials 1-160). A positive *t*-statistic indicates numeric evidence that cross-environment representational similarity is higher for the generalisable feature dimension. Statistics that cross the red dotted line indicate that the level of representational similarity is significantly higher for the generalisable feature dimension, than the non-generalisable feature dimension. Brain regions include entorhinal cortex (EC), medial orbitofrontal cortex (MOFC), hippocampus (HPC), rostral anterior cingulate cortex (RACC), lateral prefrontal cortex (LPFC), precuneus (PREC) and inferior parietal cortex (IPC).

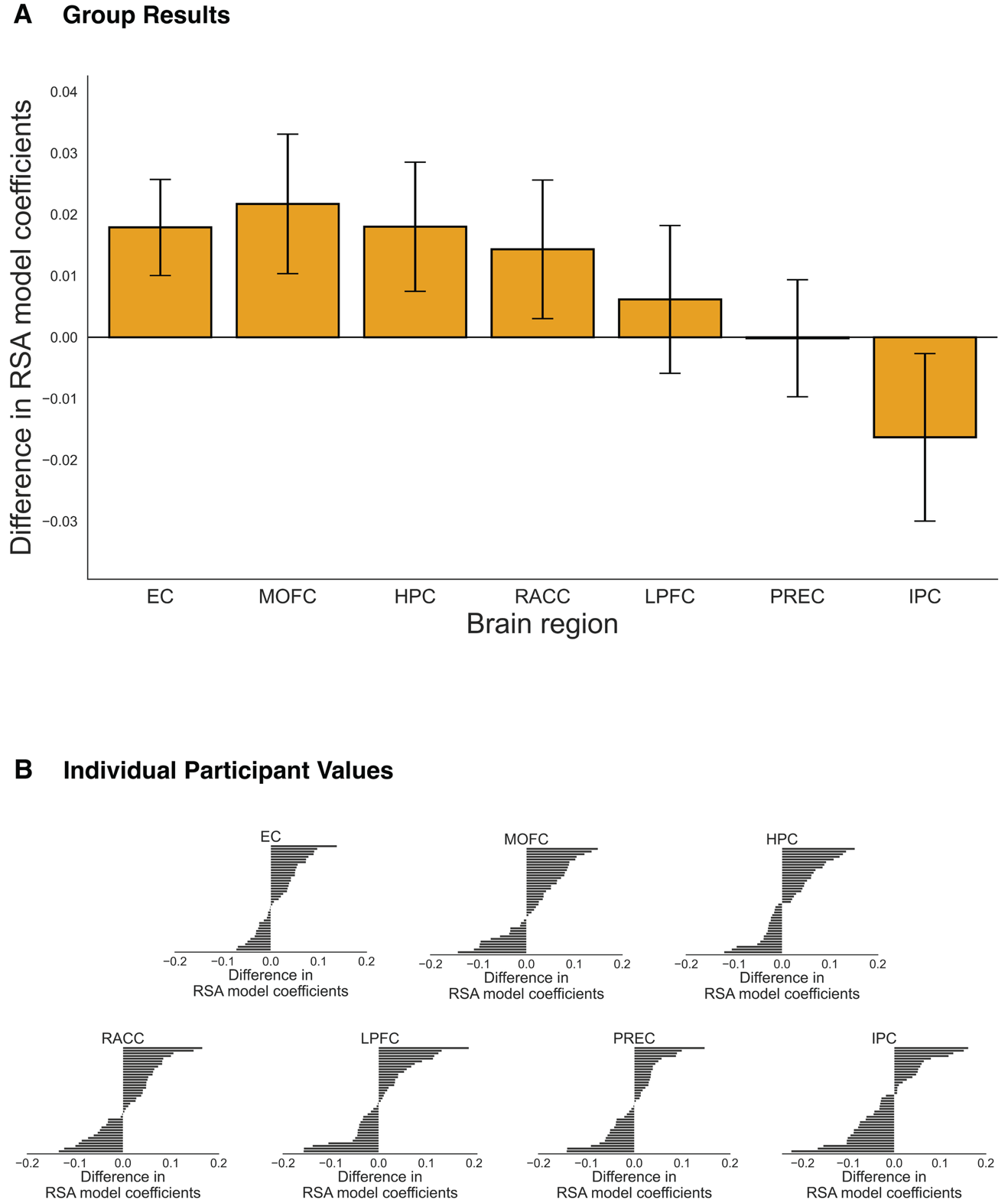

*Figure S4.* The difference between gen. and non-gen. model coefficients, for all regions of interest. Model coefficients are based on representational similarity analyses that compare neural patterns during early trials in the novel environment (1-80) to neural patterns in the familiar environment (trials 1-160). **A:** The mean difference for each region. Error bars show standard error of the mean. **B:** The neural difference for individual participants in each region. The x-axis shows the difference magnitude and each horizontal bar corresponds to an individual person. Brain regions include entorhinal cortex (EC), medial orbitofrontal cortex (MOFC), hippocampus (HPC), rostral anterior cingulate cortex (RACC), lateral prefrontal cortex (LPFC), precuneus (PREC) and inferior parietal cortex (IPC).

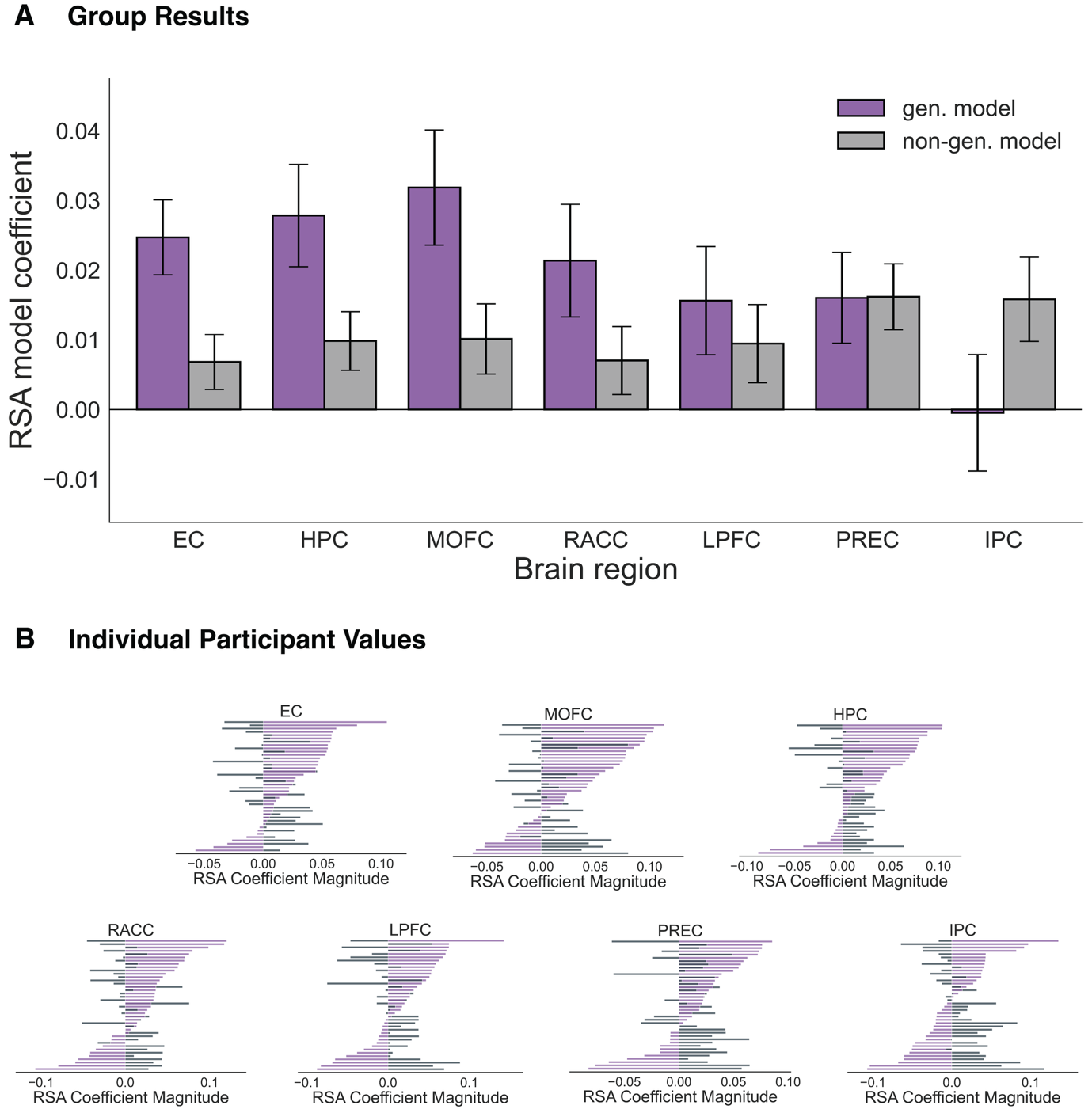

*Figure S5.* Gen. and non-gen. model coefficients for all regions of interest. Model coefficients are based on representational similarity analyses that compare neural patterns during early trials in the novel environment (1-80) to neural patterns in the familiar environment (trials 1-160). **A:** The mean coefficient magnitude for each region. Magnitudes for the gen. and non-gen. models are shown with purple and grey bars, respectively. Error bars show standard error of the mean. **B:** The coefficient magnitudes for individual participants in each region. The x-axis denotes the magnitude. Each horizontal line corresponds to an individual person, with a purple portion for their gen. model coefficient and a grey portion for their non-gen. model coefficient. Brain regions include entorhinal cortex (EC), medial orbitofrontal cortex (MOFC), hippocampus (HPC), rostral anterior cingulate cortex (RACC), lateral prefrontal cortex (LPFC), precuneus (PREC) and inferior parietal cortex (IPC).

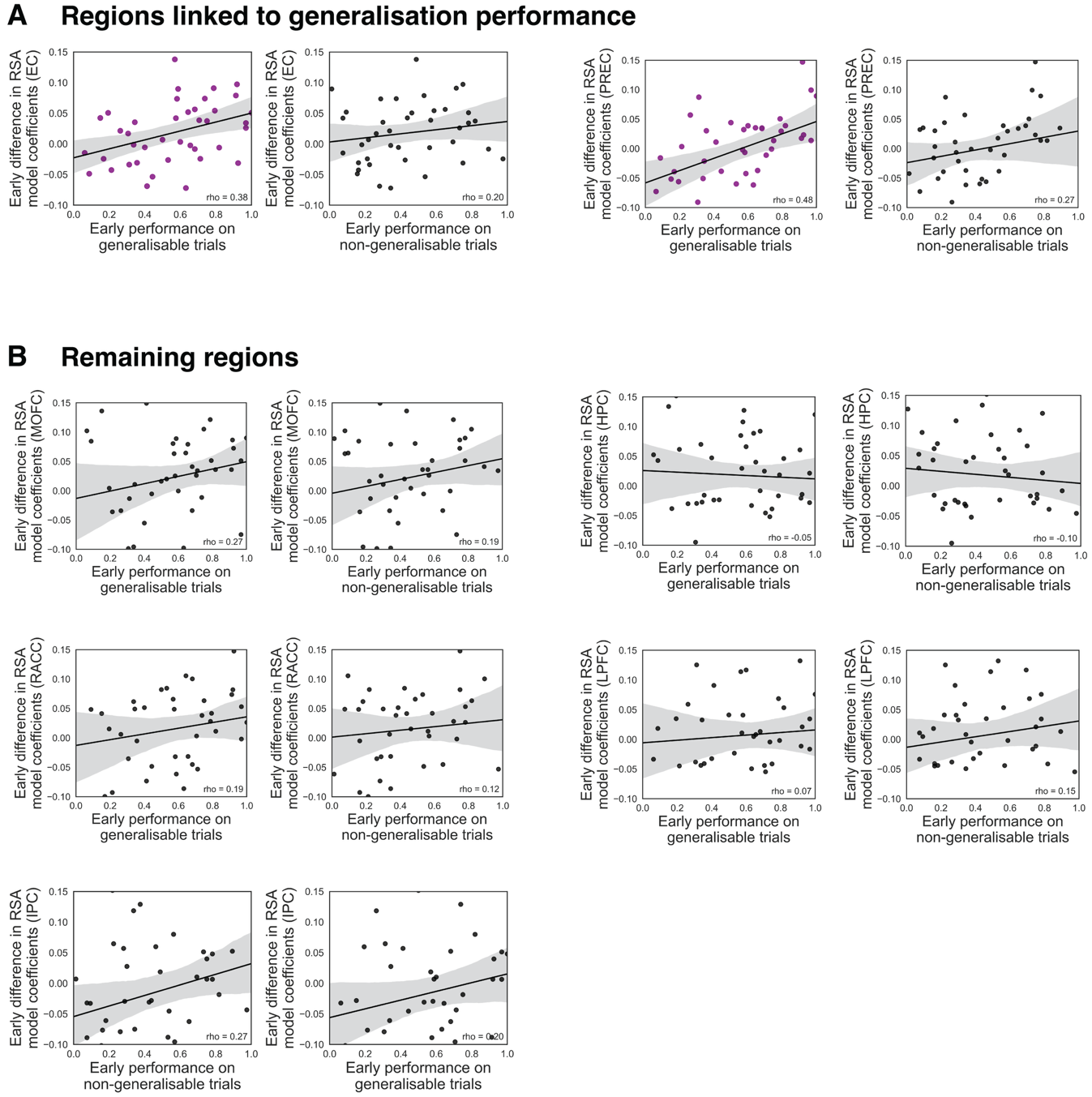

*Figure S6.* Correlations between neural signalling and generalisation performance for all regions of interest. Neural signalling refers to the difference in gen. and non-gen. model coefficients. These model coefficients are based on representational similarity analyses that compare neural patterns during early trials in the novel environment (1-80) to neural patterns in the familiar environment (trials 1-160). **A:** Brain regions with a significant Spearman correlation between neural signalling and performance during the early trials in the novel environment. This includes the entorhinal cortex (EC) and the precuneus (PREC), which show a significant correlation between early neural signalling and early performance on the generalisable blocks. These significant correlations are indicated with purple dots (EC: *ρ*(38)=0.38, *p*=0.029, PREC: *ρ*(38)=0.48, *p*=0.003). **B:** Remaining brain regions of interest, which did not show a significant correlation between neural signalling and performance (all *ρ*(38)<0.27, all *p*-values>0.19). These include medial orbitofrontal cortex (MOFC), hippocampus (HPC), rostral anterior cingulate cortex (RACC), lateral prefrontal cortex (LPFC), precuneus (PREC) and inferior parietal cortex (IPC). **A-B:** Dots show individual participant values. Black lines indicate linear fits the data and gray lines indicate 95% confidence intervals of the fits.
